# EEG Reveals Robust Within-Person but Unstable Between-Person Neural Encoding of Pain

**DOI:** 10.64898/2025.12.11.693623

**Authors:** Laura Tiemann, Felix S. Bott, Elisabeth S. May, Moritz M. Nickel, Vanessa D. Hohn, Cristina Gil Ávila, Nicolò Bruna, Paul Theo Zebhauser, Markus Ploner

## Abstract

The perception of pain varies both within and between individuals, even when sensory input remains constant. Understanding how the brain encodes these intra- and interindividual variations is essential for elucidating the neural mechanisms of pain in health and disease. Yet, previous findings have been inconsistent, and their robustness, in terms of both repeatability and replicability, has remained unclear. Here, we used electroencephalography (EEG) in 161 healthy participants to re-investigate the neural correlates of intra- and inter-individual variations in the perception of brief painful stimuli, independent of stimulus intensity. Using Bayesian multivariate multi-model regression, we related pain ratings to evoked (N1, N2, P2) and induced oscillatory responses (alpha, beta, gamma). To assess the robustness of our findings, the experiment was repeated after four weeks in the same participants and replicated in an independent cohort (*n* = 111). This design allowed us to examine within-person variability at both short (moment-to-moment) and long (day-to-day) timescales. Inter-individual differences in pain were primarily associated with the P2 response, an effect that was repeatable in the same but not replicable in the independent cohort. In contrast, intra-individual variations were explained by a multicomponent EEG pattern that was both repeatable across time and replicable across cohorts. These findings demonstrate that intra- and inter-individual variability in pain is differentially encoded in the human brain and reveal greater robustness of within-person brain-behavior associations. EEG markers may therefore be more suitable for tracking longitudinal changes in pain within individuals than for comparing pain sensitivity across individuals.

## Introduction

The experience of pain is inherently variable, both within and between individuals [1–3]. This variability arises from the dynamic interaction of sensory, cognitive, and affective processes and represents a core characteristic of pain perception in health and its alterations in chronic pain disorders. Understanding how the brain encodes these intra- and inter-individual variations is essential for advancing mechanistic models of pain and for developing reliable neural markers to guide diagnosis and treatment [2, 4].

Despite extensive research, the neural bases of this variability remain incompletely understood [5, 6]. Studies investigating inter-individual differences in pain sensitivity have yielded inconsistent findings across both functional magnetic resonance imaging (fMRI) and electroencephalography (EEG) recordings [5–12]. For example, while some EEG studies identified associations between individual pain sensitivity and induced responses at gamma frequencies [8], others identified associations with theta or combined theta-gamma responses [9, 10]. These discrepancies may reflect differences in sample size, analysis pipelines, or untested assumptions about the robustness of brain-behavior relationships.

Evidence for neural signatures of intra-individual variability of pain at short time scales (moment-to-moment / trial-to-trial) has been more consistent. Previous work has shown that brain activity in regions such as the primary somatosensory cortex, insula, anterior cingulate cortex, and prefrontal cortex tracks within-person fluctuations in pain perception across imaging modalities [6, 13–16]. In EEG studies, such short-term fluctuations have most reliably been associated with the amplitude of the N2/P2 complex [8, 17, 18] and induced responses at gamma frequencies [8, 19]. However, it remains unknown whether the neural patterns that encode short-term fluctuations also account for variability at longer time scales (day-to-day / session-to-session), even though such long-term variations might be particularly relevant in clinical contexts.

Addressing these inconsistencies and open questions requires a systematic assessment of the robustness of brain-behavior relationships. Robustness in general refers to the consistency of findings across variations in data characteristics or analytical approaches. Here, we focus on two empirical manifestations of robustness: *repeatability,* which refers to the consistency of results when the same participants complete the experiment again, and *replicability,* which refers to the consistency of results across independent cohorts. Together, these dimensions provide crucial insights into the reliability and generalizability of neurophysiological markers of within-person and between-person variability in pain perception.

In the present study, we therefore re-examined the neurophysiological signatures of within-person and between-person variability in the perception of brief painful laser stimuli and systematically evaluated their robustness by testing both repeatability and replicability. We complemented the investigation of short-term, trial-by-trial variability with the assessment of longer-term variability across sessions. To this end, we collected EEG data from 161 healthy participants who completed the same experiment twice, about four weeks apart. We related trial-level and session-level pain ratings to evoked (N1, N2, P2) and induced oscillatory (alpha, beta, gamma) responses using Bayesian multi-model regression. Finally, we assessed the replicability of our findings in an independent cohort (n = 111).

This combined design allowed us to disentangle and directly compare the robustness of within-person versus between-person neural encodings of pain. Our findings reveal that while inter-individual associations are limited in their replicability, intraindividual associations are robust across time and cohorts. Together, these findings highlight the potential of within-person EEG markers for the longitudinal tracking of pain.

## Results

### Overview

In the present study, we investigated how brain responses to noxious stimuli relate to inter-individual and intra-individual variability in pain perception (Fig. 1A). Intra-individual variability was assessed at both short (moment-to-moment) and long (day-to-day) timescales. The experiment consisted of two sessions approximately one month apart. In each session (Fig. 1B), participants received 80 brief painful stimuli to the dorsum of the left hand and provided verbal pain ratings for each stimulus. Stimuli were delivered at four different intensities. To optimize the signal-to-noise ratio, we only included trials with stimuli of the two highest intensities.

**Figure 1:**
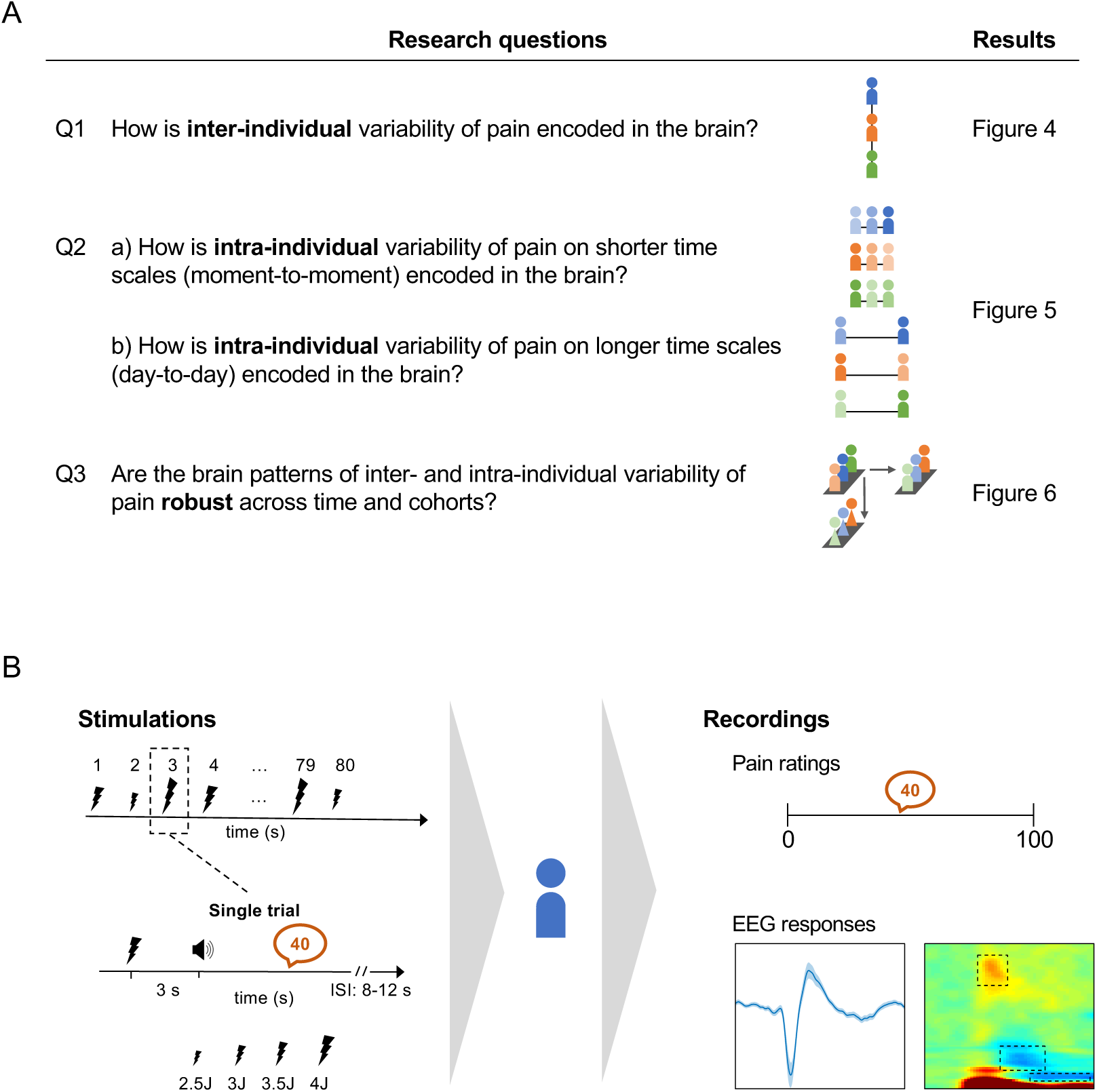
Study overview. A) Research questions and corresponding result figures. B) During the experiment, participants received 80 brief painful laser stimuli to the dorsum of their left hand at four different fixed intensities (2.5, 3.0, 3.5, 4.0 J). An auditory cue prompted participants to verbally rate their pain intensity on a numerical rating scale ranging from 0 (“no pain”) to 100 (“maximally tolerable pain”). Concurrently, EEG was recorded with a 64-channel system. Throughout the experiment, participants sat in a comfortable chair with eyes closed, wore protective goggles, and listened to white noise via headphones to mask ambient sounds.

Brain responses were quantified using EEG. We extracted six commonly analyzed features of event-related brain activity: the amplitudes of the N1, N2, and P2 evoked responses, and the power of induced oscillatory responses at alpha, beta, and gamma frequencies. Pain perception was measured using verbal pain ratings on a numerical rating scale ranging from 0 (no pain) to 100 (maximally tolerable pain).

To quantify associations between brain responses and pain ratings, we used Bayesian multi-model regression, treating the six EEG features as predictors and the pain ratings as the outcome variable. Bayesian multi-model inference enabled variable selection and parameter estimation within a unified framework by averaging over models with all plausible combinations of predictors. This approach accounts for model uncertainty and yields more stable and less biased inferences about relationships between brain responses and pain ratings [20].

The study and its analysis were preregistered at ClinicalTrials.gov (https://clinicaltrials.gov/study/NCT05616091) and the OSF Registries (https://osf.io/wfjhq).

### Brain responses to noxious stimuli

To confirm that the EEG responses in the present study reproduced the well-established patterns elicited by brief noxious laser stimuli, we computed grand averages of evoked potentials and time-frequency representations (Fig. 2). Consistent with previous work [8, 21], the evoked potential at the Cz-Avg montage showed pronounced negative and positive deflections at approximately 240 ms and 375 ms post stimulus, respectively (Fig. 2A). In addition, in line with prior findings [8, 21, 22], alpha- and beta-band oscillatory activity decreased around 700 ms and 350 ms, respectively, whereas gamma oscillatory activity increased around 250 ms post-stimulus (Fig. 2B). Together, these results confirm that the recorded EEG activity closely reflects the characteristic pattern of EEG responses to this type of noxious stimuli.

**Figure 2:**
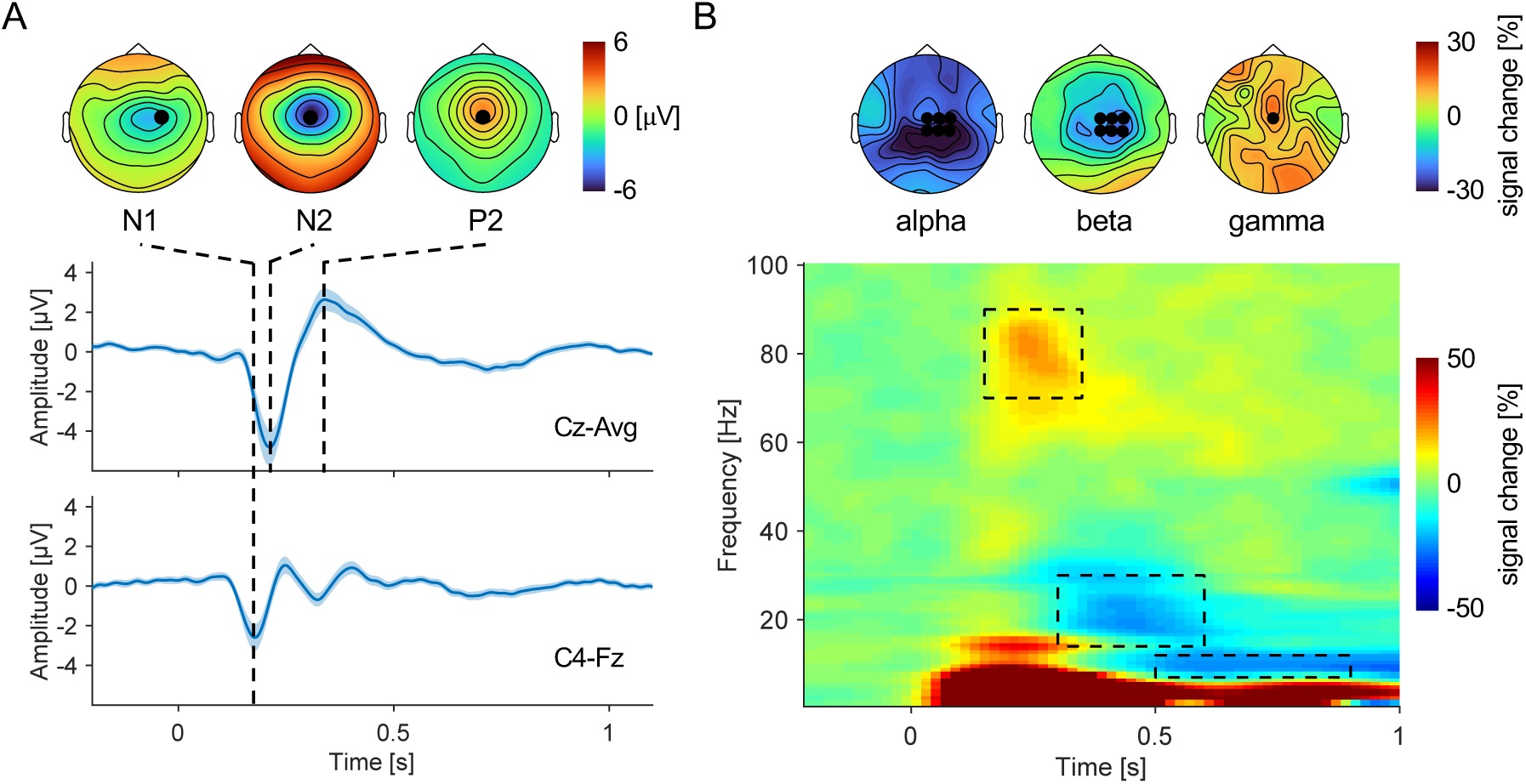
Brain responses to noxious stimuli. A) Grand average of event-related potentials for the Cz-Avg montage (to visualize the N2 and P2 responses) and the C4-Fz montage (to visualize the N1 response). The shaded blue bands represent +/- 2 SEM. Scalp topographies show the spatial distributions of voltage for the respective montages. \ Positions of measurement electrodes are marked as black dots. B) Grand average time-frequency representation (TFR) showing changes in oscillatory power relative to a pre-stimulus baseline [-1 to 0 s]. For visualization, TFRs at Cz are presented. Scalp topographies illustrate the spatial distributions of relative power changes, averaged within the indicated time-frequency windows. Measurement electrode included later for the quantification of oscillatory power in the different frequency bands are marked as black dots. The grand averages displayed in A) and B) represent data from the first of two internal recording sessions. Corresponding visualizations for the second internal session and for the externally acquired data are provided in the supplementary materials (Fig. S1).

### Brain patterns of *inter-individual* variability of pain

To assess brain patterns of inter-individual variability of pain, we first quantified individual pain ratings and corresponding EEG responses. Individual pain ratings were obtained by averaging single-trial pain ratings across trials and stimulus intensities within each participant. Individual brain responses were computed analogously by averaging evoked potentials and time-frequency representations within each participant. The distribution of individual pain ratings is shown in Fig. 3A. Corresponding distributions of brain responses are provided in the supplementary materials (Fig. S3 and S4).

**Figure 3:**
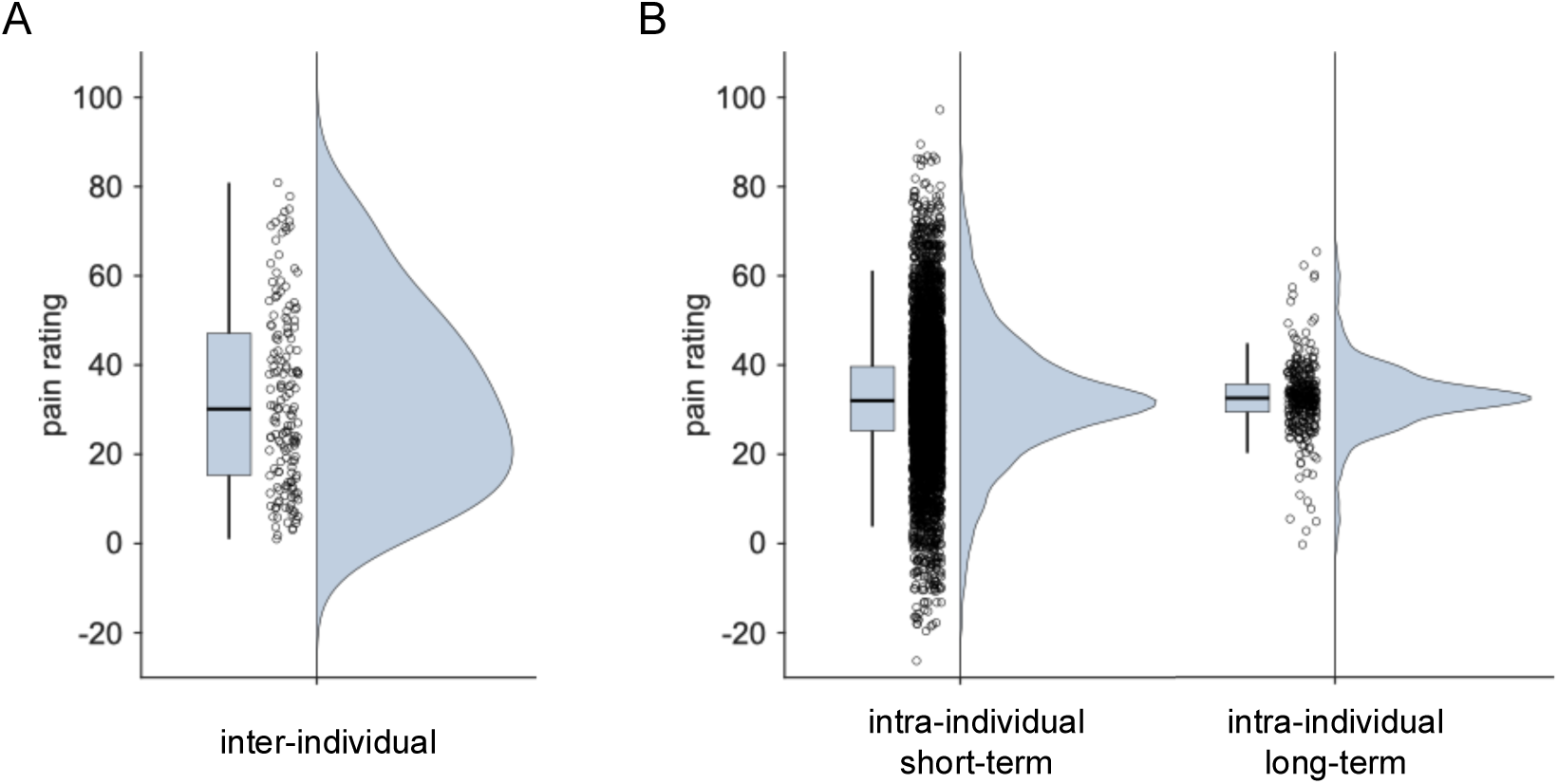
Pain ratings. A) Individual pain ratings reflecting inter-individual pain variability were obtained by averaging single-trail pain ratings for each individual, first within each stimulus intensity and then across stimulus intensities, yielding one rating value per individual. B) Trial-level pain ratings (left) reflecting intra-individual short-term pain variability were obtained by computing the means of single-trial pain ratings for each individual and stimulus intensity, and then subtracting these means from the corresponding single-trail pain ratings, yielding one centred rating value per trial. Session-level pain ratings (right) reflecting intra-individual pain variability at longer timescales were obtained by combining and then averaging individual pain ratings from session 1 and session 2 for each individual, and then subtracting these means from the corresponding individual pain ratings, yielding two centred rating values per individual. For visualization only, trial-wise and session-wise pain ratings were re-centred to the mean of the inter-individual distribution. The presented pain ratings originate from the first (first and second in the case of session-wise ratings) of two internal recording sessions. Pain rating distributions for the second internal session and for the externally acquired data are provided in the supplementary materials (Fig. S2).

The Bayesian multi-model analysis yielded strong evidence for models relating EEG responses to pain perception across individuals (Fig. 4, BF_10_ > 200 for the top 5 models; indicating that the evidence for these models is more than 200 times stronger than for the null model). To control for potential confounding effects of age and gender, both variables were included in every model, including the null model. The best model comprised only the P2 amplitude as a predictor and showed the largest change from prior to posterior model odds (BF_M_ = 25). Notably, all of the top 5 models also included the P2 amplitude. Across these models, the proportion of variance explained in pain ratings ranged from R^2^ = 0.14 to R^2^ = 0.17.

**Figure 4:**
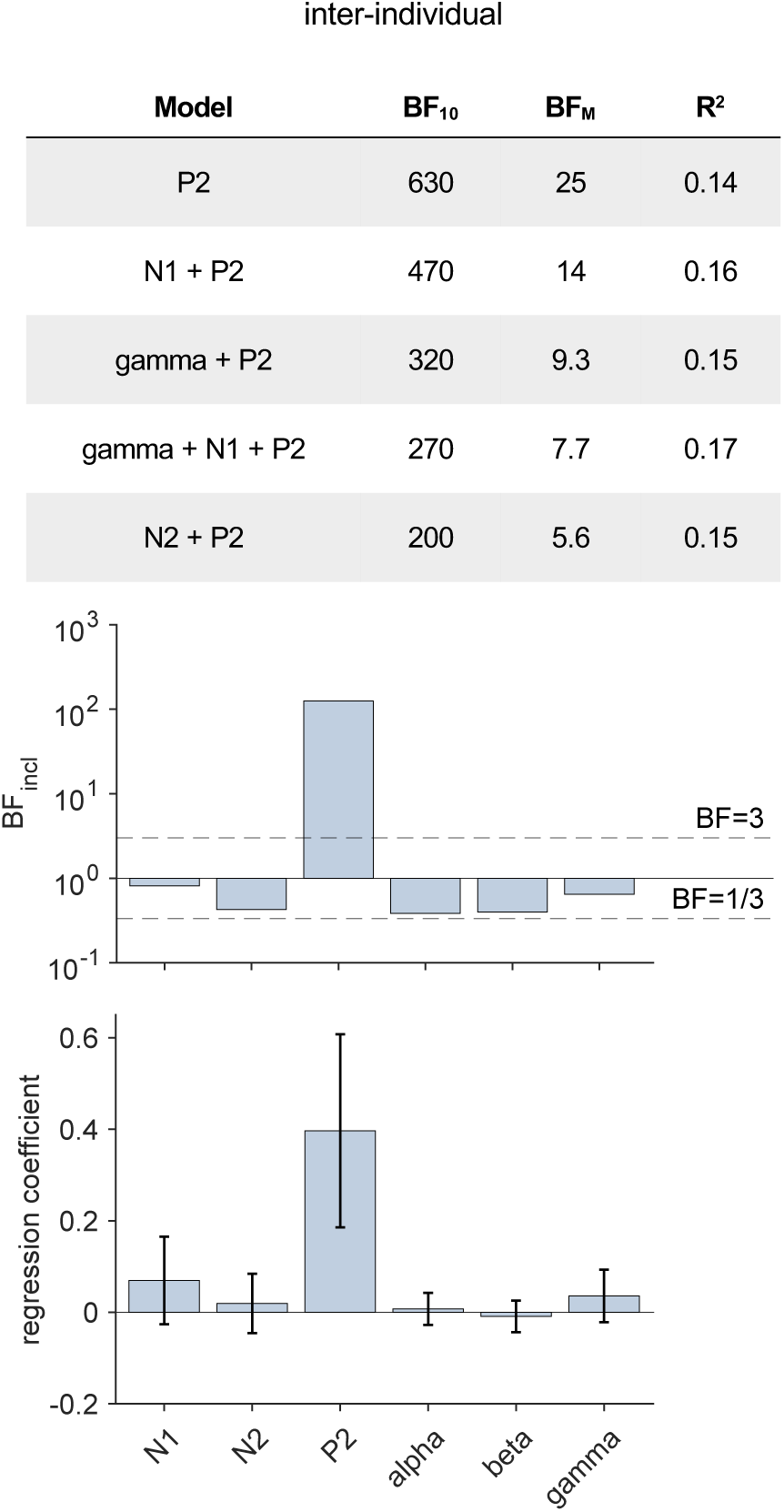
Bayesian multi-model analysis of inter-individual variability. Top: The five models with the highest evidence relative to the null model (BF_10_) are shown, along with their ratios of prior and posterior model odds (BF_M_) and their explained variance (R^2^). Middle: Inclusion Bayes Factors (BF_incl_), representing the evidence for including each predictor averaged across all models considered. Bottom: Model-averaged posterior mean estimates for each predictor coefficient. Error bars indicate +/-2 standard deviations of the model-averaged posterior coefficient distribution.

Model-averaged estimates further confirmed the dominant role of the P2 amplitude in explaining inter-individual variability in pain (BF_incl_ = 122; indicating that the data are 122 times more likely under models including this predictor than under models excluding it). Posterior coefficient estimates indicated a positive association between P2 amplitude and individual pain sensitivity. For all other EEG responses, analyses provided anecdotal evidence against their contribution (1/3 < BF_incl_ 1/3 < 1).

Together, these findings demonstrate that EEG responses to noxious stimuli, and particularly the P2 amplitude, explain inter-individual differences in pain perception.

### Brain patterns of *intra-individual* variability of pain

To examine intra-individual variability at short timescales, we analyzed single-trial pain ratings together with their corresponding brain responses. To isolate fluctuations in pain perception that were independent of stimulus intensity, we centred all variables separately within each participant-intensity pair; that is, we subtracted each participant’s average rating for a given intensity from their trial-wise ratings. The distribution of centred single-trial pain ratings is shown in the left panel of Fig. 3B. Corresponding distributions of brain responses are provided in the supplementary materials (Fig. S3 and S4).

The Bayesian multi-model analysis yielded strong evidence for models relating brain responses to pain ratings within individuals at short timescales (Figure 5A, BF_10_ > 10^59^for the top 5 models). The best model comprised all six brain responses and showed the largest change from prior to posterior model odds (BF_M_ = 80). Across the top five models, the proportion of variance explained in pain ratings was approximately R^2^ ≈ 0.08. Model-averaged estimates confirmed that all six brain responses contributed to explaining short-term, intra-individual variability (BF_incl_ > 10 for all brain responses), with the strongest evidence observed for the N2 amplitude (BF_incl_ > 10^45^). For all EEG responses, stronger responses (more negative decrease / more positive increase) were associated with higher pain ratings. Together, these findings demonstrate that all six EEG responses to noxious stimuli contribute to intra-individual variability in pain perception at short timescales.

**Figure 5:**
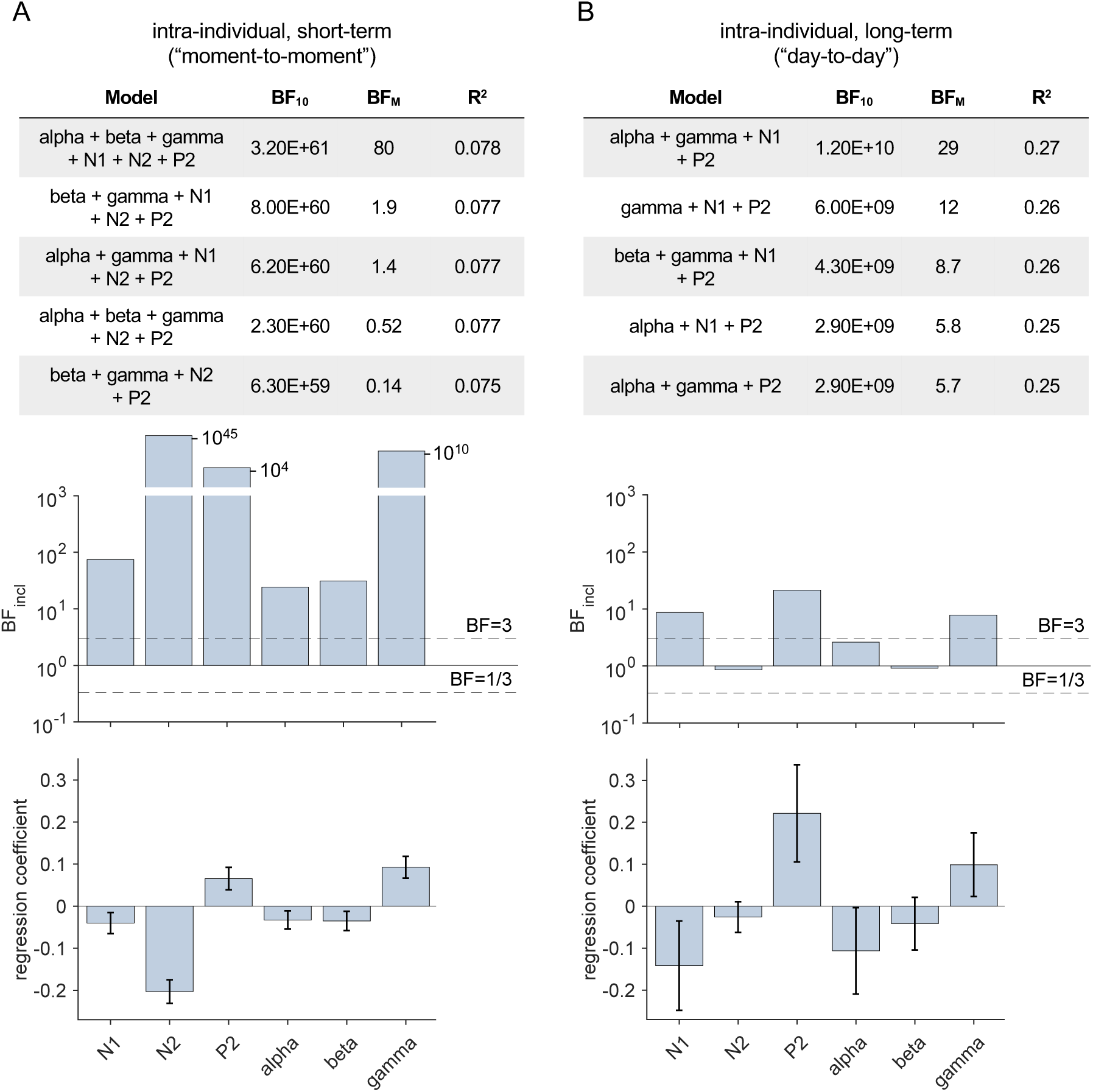
Bayesian multi-model analysis of intra-individual variability. A) moment-to-moment variability. B) day-to-day variability. A) + B) Top: The five models with the highest evidence relative to the null model (BF_10_) are shown, along with their ratios of prior and posterior model odds (BF_M_) and their explained variance (R^2^). Middle: Inclusion Bayes Factors (BF_incl_), representing the evidence for including each predictor averaged across all models considered. Bottom: Model-averaged posterior mean estimates for each predictor coefficient. Error bars indicate +/-2 standard deviations of the model-averaged posterior coefficient distribution.

To assess intra-individual variability at longer timescales, we combined pain ratings and brain responses from sessions 1 and 2, yielding two data points per participant. As in the short-timescale analysis, we centred all variables within each participant to remove between-subject variability. The distribution of centred, session-level pain ratings is shown on the right in Fig. 3B. Corresponding distributions of brain responses are provided in the supplementary materials (Fig. S3 and S4).

The Bayesian multi-model analysis again yielded strong evidence for models relating brain responses to pain ratings within individuals at longer timescales (Figure 5B, BF_10_ > 10^9^ for the top 5 models). The best model comprised the alpha, beta, N1, and P2 responses and showed the largest change from prior to posterior model odds (BF_M_ = 29). The proportion of variance explained by the top five models ranged from R^2^ = 0.25 to R^2^ = 0.27. Model-averaged estimators indicated that, in particular, the N1, P2, and gamma responses contributed to explaining longer-term intra-individual variability of pain perception (BF_incl_ > 10 for P2 amplitude, BF_incl_ > 3 for N1 amplitude and gamma power). As in the short-timescale analysis, stronger brain responses were associated with higher pain ratings. Together, these findings indicate that multiple EEG responses, particularly the P2, N1, and gamma responses, contribute to intra-individual variability in pain perception at longer time scales. Moreover, the directions of the associations were consistent across short and long timescales, indicating stable within-individual brain-behavior relationships over time.

### Robustness

Next, we assessed the robustness of brain response patterns of inter- and intra-individual variability of pain. Robustness was evaluated in terms of repeatability (consistency across the two sessions in the same cohort) and replicability (consistency in an independent cohort). For both analyses, we estimated model-averaged coefficients in one dataset, applied them to predict pain ratings in another dataset, and quantified the association between predicted and observed pain ratings. To test repeatability, coefficients were estimated from internal session 1 to predict pain ratings of internal session 2, and vice versa. To test replicability, coefficients estimated from either internal session were applied to an external dataset, and vice versa. The external dataset comprised EEG recordings of healthy individuals who received brief painful laser stimuli to the left hand at two individually adjusted intensities across 30 trials. To optimize the signal-to-noise ratio, we only included the 15 higher-intensity trials and excluded participants with fewer than 5 clean trials, yielding a final sample of 111 individuals. As the external dataset comprised only a single session, the replicability of brain response patterns explaining session-to-session variability of pain perception was not assessed.

As shown in Figure 6A, we found strong evidence for the repeatability of brain response patterns explaining inter-individual variability in pain (BF > 10^3^, R^2^ > 0.11). Coefficients estimated in session 1 generalized to session 2, and vice versa. In contrast, we found evidence against the replicability of these patterns in the independent cohort. Coefficients estimated from session 1 or session 2 provided evidence against replicability when applied to the external dataset (BF < 1/3, R^2^ ≈ 0). Conversely, coefficients estimated from the external dataset provided evidence against replicability when applied to the internal session 1 or session 2 datasets (BF < 1/3, R^2^ ≈ 0). Results of the Bayesian multi-model analysis of inter-individual variability in the external dataset are provided in the supplementary materials (Fig. S5 A).

**Figure 6:**
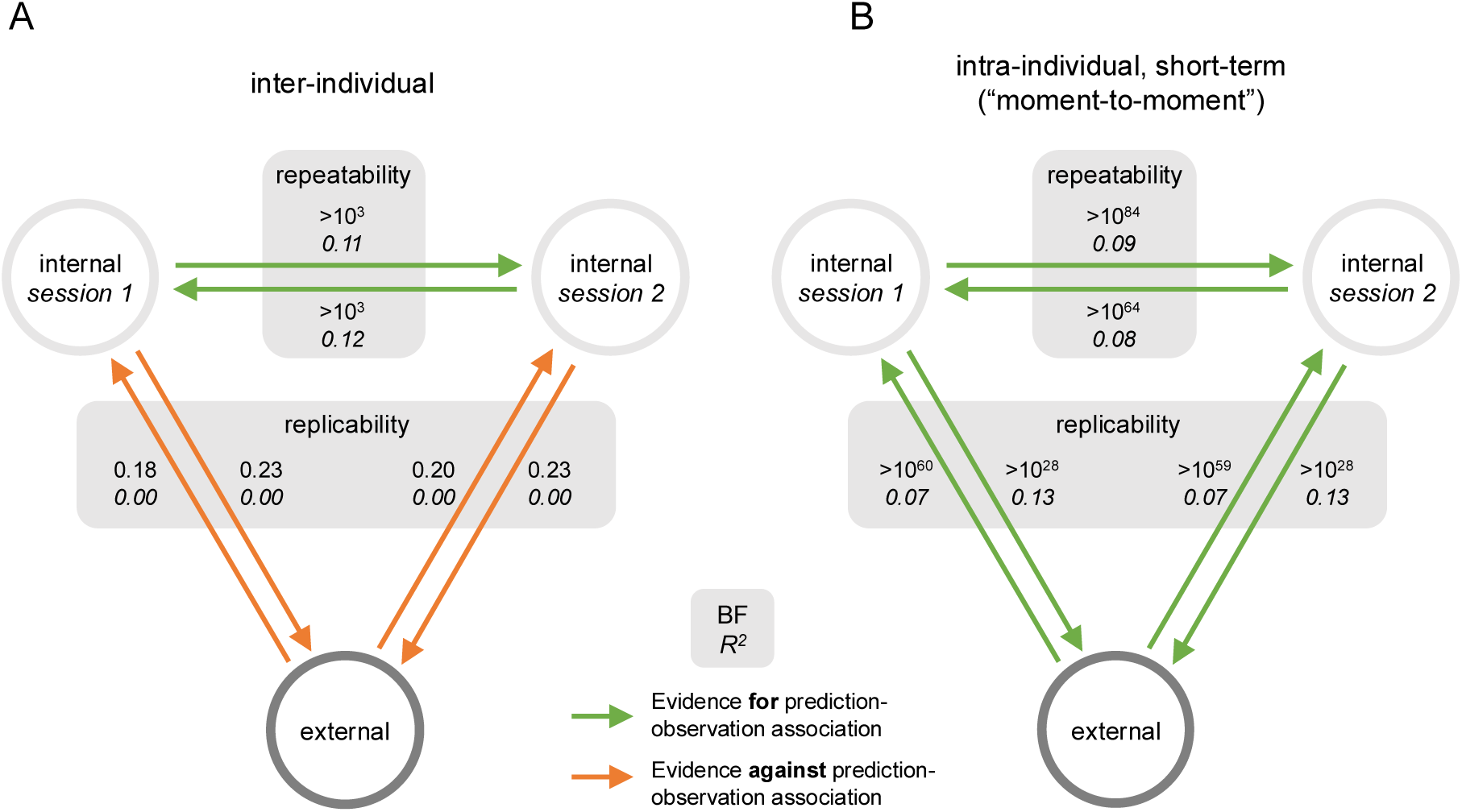
Robustness of the neural patterns of the inter- and intra-individual variability of pain. A) Robustness of patterns associated with inter-individual variability. B) Robustness of patterns associated with intra-individual moment-to-moment variability. A) + B) Each circle represents one dataset. The two internal datasets share the same participant cohort across sessions, while the external dataset includes an independent cohort. Shaded regions illustrate repeatability (consistent results when repeating the experiment in the same cohort) and replicability (consistent results when repeating a similar experiment in a different cohort). To test repeatability and replicability, model-averaged coefficients were estimated from one dataset (“training set”) and then used to predict pain ratings in another (“test set”). Arrow tails and heads denote training and test sets, respectively. Numbers beside arrows represent BFs quantifying evidence for an association between predicted and observed pain ratings, along with the R^2^-score, i.e., the proportion of variance in observed pain ratings explained by the predictions.

As shown in Figure 6B, we found strong evidence for the repeatability of brain response patterns explaining short-term intra-individual variability (BF > 10^64^, R^2^ > 0.08). Coefficients estimated in session 1 generalized to session 2, and vice versa.

Crucially, these patterns were also replicable in the independent cohort. Coefficients estimated from session 1 or session 2 provided strong evidence for replicability when applied to the external dataset (BF > 10^28^, R^2^ ≈ 0.13). Conversely, coefficients estimated from the external dataset generalized to both internal sessions (BF > 10^59^, R^2^ = 0.07). Results of the Bayesian multi-model analysis for the external dataset are provided in the supplementary materials (Fig. S5 B).

## Discussion

In this study, we investigated how variability in EEG responses to noxious stimuli relates to variability in pain perception and how robust these associations are (Fig. 7). We found strong evidence for brain-behavior relationships at both the inter- and intra-individual levels. Inter-individual variability in pain was primarily explained by a single EEG response, i.e., the P2 amplitude. This association was repeatable within the same cohort but did not replicate in an independent cohort. In contrast, intra-individual variability in pain was explained by a broader set of brain responses. At short timescales (moment-to-moment), all six EEG responses contributed to within-person fluctuations of pain, and this multicomponent pattern was both repeatable and replicable. At longer timescales (day-to-day), intra-individual variability was mainly associated with the P2, N1, and gamma responses. The direction of these intra-individual brain-behavior associations was consistent across both short and long timescales.

**Figure 7:**
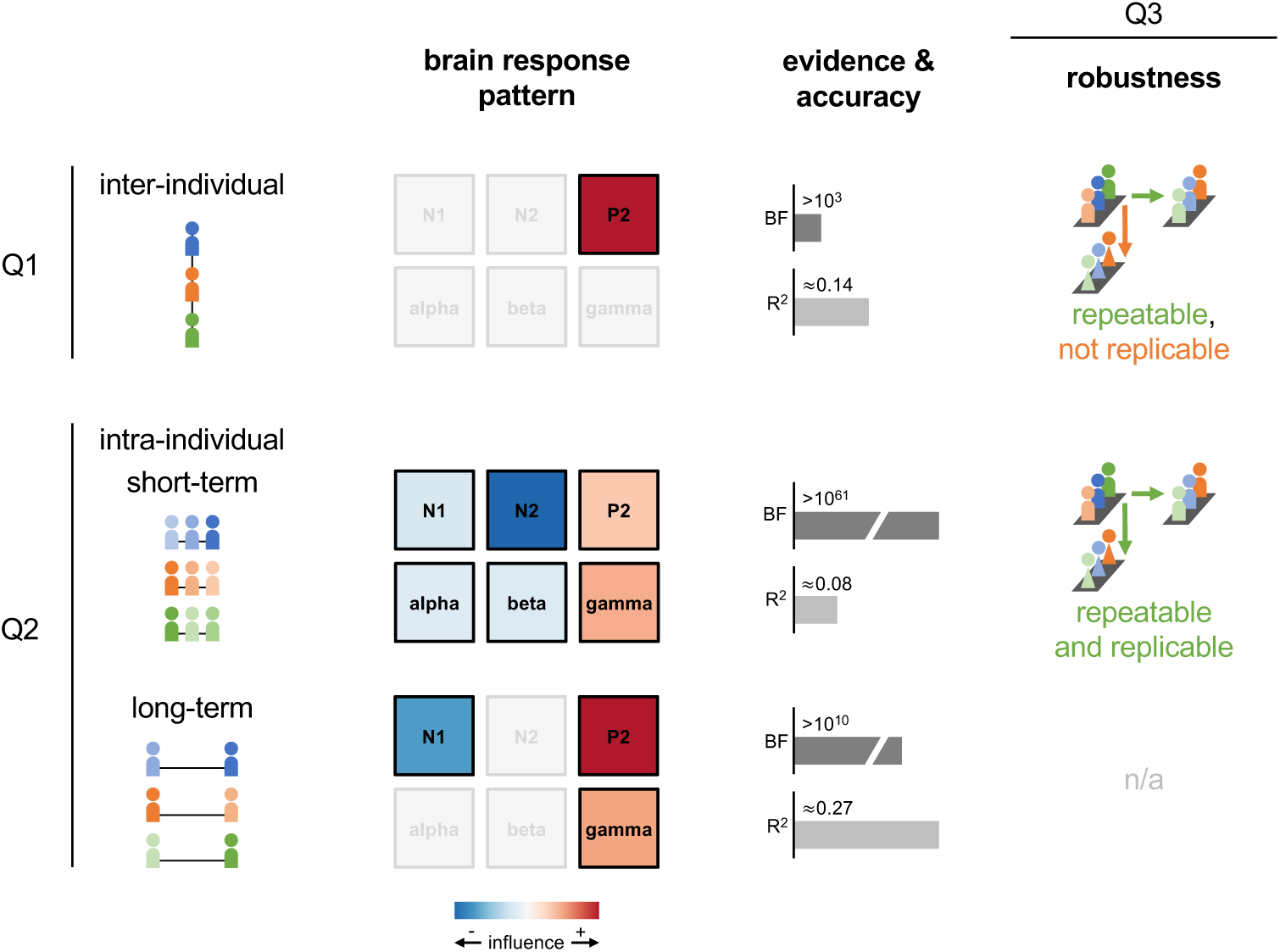
Visual summary of results. This study examined EEG response patterns underlying inter- and intra-individual variability of pain perception. Tile plots show color-coded, model-averaged coefficient estimates, indicating the magnitude and direction of influence of each EEG response on pain perception. EEG responses with BF_incl_ < 3 are shown in grey. Bar plots summarize the evidence (BF_10_) and accuracy (R^2^) of the best models for each variability component. Q1) EEG responses, and particularly the P2 amplitude, can explain inter-individual differences in pain sensitivity. Q2) EEG responses explain both short and long-term intra-individual variability of pain. To isolate variability unrelated to stimulus intensity, all variables were centred per participant and intensity in this analysis. All brain responses explained short-term variability of pain, while long-term variability of pain was predominantly explained by P2, N1, and gamma responses. Q3) EEG response patterns of inter-individual variability were repeatable but not replicable. EEG response patterns of short-term intra-individual pain variability were both repeatable within the same cohort and replicable in an independent cohort. Repeatability and replicability of brain response patterns of long-term pain variability could not be evaluated with the available dataset. Together, these findings indicate distinct EEG response patterns for inter- and intra-individual variability of pain, and highlight differences in terms of their robustness with patterns of intra-individual variability being more robust than those of inter-individual differences.

Overall, our findings indicate that distinct neural patterns underlie inter- and intra-individual variability in pain and that these patterns differ significantly in their robustness. Whereas previous studies reported that oscillatory induced responses at theta or gamma frequencies encode inter-individual differences in pain perception [8–10], we found only non-replicable evidence for a role of the P2 evoked response in accounting for such differences. In contrast, the observation that all considered EEG responses contributed to intra-individual variability at short timescales aligns well with prior work [8, 17–19]. Complementing and extending these findings, our results reveal that similar associations are also present at longer timescales, demonstrating that within-person brain-behavior relationships, at least partially, generalize across moments and days.

The absence of replicable neural patterns explaining inter-individual variability in pain may have several explanations. First, cross-sectional analyses may not appropriately capture inter-individual differences in the neural encoding of pain. However, evidence from other studies [11, 12, 23, 24], as well as the repeatability within the same cohort of the inter-individual brain patterns identified here, argues against an inherent limitation of capturing inter-individual differences in pain perception. Second, EEG may not capture the relevant neural patterns sufficiently well due to its limited spatial resolution and limited capacity to detect activity in deeper regions such as the limbic network [25]. Third, replicability may have been limited by differences between datasets in participant characteristics and experimental procedures. The two cohorts differed in age and possibly also in sociocultural factors. Additionally, the external dataset contained fewer than half as many trials per participant as the internal dataset, which likely reduced the signal-to-noise ratio. This interpretation cannot be ruled out. Fourth, pain is a complex and multidimensional experience, and its neural representation is likely equally complex. Here, we focused on canonical EEG responses, but it is possible that other EEG features (e.g., functional connectivity) or modeling techniques (e.g., deep learning) may more robustly capture inter-individual differences in pain perception.

The evidence supporting EEG response patterns of intra-individual variability in pain was compelling. Nevertheless, the proportion of variance explained in trial-level pain ratings was modest. This likely reflects two factors: First, inherent inter-individual differences in the neural encoding of pain are not captured by a group-level model. Our analysis of intra-individual variability focused on fixed effects, i.e., brain-pain relationships assumed to generalize across individuals, and therefore cannot account for person-specific deviations from this pattern. Second, the signal-to-noise ratio of single-trial measures, both in EEG features and pain ratings, is comparatively low. This interpretation is supported by the observation that explained variance increased substantially when analyzing session-level pain ratings, which aggregate information across many trials and thus improve the signal-to-noise ratio. Interestingly, models predicting session-level pain ratings explained a larger fraction of variance than models predicting trial-level pain ratings, despite the absolute variance of session-level ratings being smaller.

Our findings have broader implications for understanding how pain is represented in the brain and for developing clinically useful biomarkers. First, we analyzed inter-individual variability reflecting the goal to identify a neural pattern that can predict pain sensitivity in previously unseen individuals. Such patterns might serve different clinical purposes such as diagnostic or prognostic biomarkers and, on a conceptual level, correspond to a nomothetic, i.e, population-based approach [26]. In our analyses, this approach yielded models that explained only a modest fraction of inter-individual variance in pain ratings and were not replicable in an independent cohort. Given that our experimental paradigm already provides an optimized and highly controlled setting for probing neural representations of pain, these findings challenge the feasibility of developing robust, EEG-based diagnostic markers for clinical pain conditions.

Second, we analyzed intra-individual variability of pain reflecting the goal to identify a neural pattern that can track changes in pain perception within single individuals over time. Such patterns might serve clinical purposes as monitoring biomarkers and, on a conceptual level, largely correspond to a nomothetic approach, albeit with idiographic, i.e., individual-specific components in the form of random intercepts [23, 26, 27]. Here, this approach gave rise to models that were reproducible in an independent dataset and explained a larger fraction of variance in pain ratings at longer timescales. This suggests that more idiographic, personalized modelling approaches may better capture the neural underpinnings of variability in pain perception. Although such models do not generalize to the entire population, they may serve as valuable tools for tracking individual pain mechanisms and informing personalized pain management.

Our study has limitations, particularly regarding the specificity of the reported effects. First, it remains unclear whether our findings are specific to pain perception or extend to other sensory modalities. Second, our experimental design does not rule out the possibility that the observed EEG patterns partly reflect variability in peripheral rather than central nociceptive processing. Although questions of pain specificity and brain specificity are important in their own right, they were not the primary focus of the present study. Rather than inferring specific neural mechanisms, our goal was to characterize statistical associations between EEG measures and different components of variability in pain perception. These associations may provide a foundation for future mechanistic studies and inform the development of clinically relevant biomarkers.

In conclusion, this study provides a comprehensive characterization of how neural responses to noxious stimuli relate to pain perception. We examined variability in pain both between individuals and within individuals across short and long timescales. We rigorously assessed the robustness of our findings using a novel analytical framework in combination with a unique constellation of multiple datasets. While we found strong evidence for repeatable inter-individual associations between brain responses and pain ratings, these associations were not replicable in an independent cohort. In contrast, intra-individual associations were robust. They were repeatable, replicable, and evident across different timescales. These findings highlight the potential of individualized, longitudinal approaches for understanding the brain mechanisms of pain. They suggest that within-person EEG markers may offer a promising foundation for developing clinically useful biomarkers that can inform personalized pain management.

## Methods

### Dataset

The study uses a dataset recorded by our research group (https://www.painlabmunich.de/) at the Technical University of Munich (TUM) in Germany between December 2019 and December 2022 (https://osf.io/z2h86/). We refer to this dataset as Set_Munich. The study protocol was approved by the Ethics Committee of the Medical Faculty of the TUM and conducted following the latest version of the Declaration of Helsinki. The study was preregistered at ClinicalTrials.gov (https://clinicaltrials.gov/study/NCT05616091) prior to data collection. The present analysis and hypotheses were preregistered on the OSF Registries (https://osf.io/wfjhq) prior to conducting any analyses related to this study’s research questions. While parts of the dataset have been previously analyzed in a separate project [28] addressing different research questions (https://osf.io/s75q6/), the authors had no prior knowledge of the effects investigated here.

### Participants

We used non-probability sampling to recruit equal numbers of female and male participants, as well as equal numbers of individuals aged ≤ 40 years and > 40 years. Recruitment was conducted via advertisements posted across the Technical University of Munich (TUM), its university hospital, and their associated websites. In addition, individuals from previous studies who had provided consent to be recontacted were invited to participate. This approach yielded a sample with a broad age range but predominantly consisting of individuals who are read as white and are pursuing or holding higher education degrees. Therefore, the study sample represents a specific population and is not necessarily representative of Germany or any particular region.

The present data analysis includes all participants for whom experimental pain recordings were available for both sessions. Of the total sample of n = 166, four participants did not return for the second session. Of the participants who returned for the second session, one was excluded due to not having a sufficient number of artifact-free trials after preprocessing (see below for more information on this criterion). Thus, the final sample size was n = 161 (82/80 female/male; age = 40 years ± 18 years (mean ± SD), range 18 to 86 years). Inclusion criteria were age ≥ 18 years and right-handedness. Right-handed participants were selected to avoid potential variability in pain sensitivity related to stimulation of the dominant versus non-dominant hand [29, 30]. Exclusion criteria included pregnancy; neurological or psychiatric disorders (e.g., epilepsy, stroke, depression, anxiety disorders); severe systemic illnesses (e.g., cancer, diabetes); dermatological conditions (e.g., dermatitis, psoriasis, eczema); current or recurrent pain; regular or recent use of centrally acting, antibiotic, or analgesic medication; previous neurosurgical head procedures; history of head trauma with loss of consciousness; a history of fainting spells or syncope; and adverse reactions to prior electrical, magnetic, or thermal stimulation.

### Procedure and paradigm

Participants completed two identical sessions on separate days, approximately one month apart. For premenopausal female participants, both sessions were scheduled between days 5 and 10 of the follicular phase to control for potential hormonal influences on pain perception and related neuronal processes [31–34]. For 155 participants, the interval between sessions was 30 ± 9 days (mean ± SD), ranging from 17 to 98 days. For the remaining 7 participants, the interval exceeded 200 days (292 ± 89 days, range: 223–478 days) due to recording delays caused by the COVID-19 pandemic. The experiment was conducted in German.

At the beginning of each session, participants completed clinical and demographic questionnaires, including assessments of sleep quality, anxiety and depression, affect, and pain sensitivity. These data are not analyzed in the present study. Next, participants were familiarized with the laser stimulation and rating procedure. To this end, 20 laser stimuli were applied in a pseudo-randomized sequence, with five stimuli at each of the four intensities used later in the main experiment. Subsequently, EEG caps were prepared.

Each recording session began with a five-minute eyes-closed resting-state EEG, during which participants were instructed to remain relaxed but awake. These data are not analyzed here. Immediately afterwards, participants completed the experimental pain paradigm, which consisted of 80 brief laser stimuli at four different intensities applied to the dorsum of the left hand, again with eyes closed. As defined in the preregistration, we only included trials with stimuli of the two highest intensities in the present analysis. Three seconds after each stimulus, an auditory cue prompted participants to verbally rate the perceived pain intensity on a numerical scale from 0 (“no pain”) to 100 (“maximally tolerable pain”). During all recordings, participants sat in a comfortable chair, wore protective goggles, and listened to white noise via headphones to mask ambient sounds.

For further details on the data collection procedure, including the acquisition of demographic and psychological variables and skin conductance responses, which are also not analyzed here,please refer to the study’s preregistration (https://clinicaltrials.gov/ct2/show/NCT05616091). These additional variables will be addressed in a separate manuscript.

### Noxious stimulation

Pain stimuli were delivered to the dorsum of the left hand using a laser device (DEKA Stimul 1340, Calenzano, Italy) with a wavelength of 1340 nm, a pulse duration of 4 ms, and a spot diameter of 7 mm. Stimuli were administered in a pseudo-randomized order at four fixed intensities (2.5, 3, 3.5, and 4 J; 20 stimuli per intensity) with interstimulus intervals ranging from 8 to 12 s. These intensity levels were selected based on previous studies using the same device and parameters [8, 14, 15, 35], as well as internal piloting. To keep the objective stimulus intensities/inputs constant between participants, stimulation intensities were fixed [36]. Unbeknownst to the participant, stimuli were applied in 4 blocks of 20 stimuli, with 5 stimuli at each intensity. Within each block, stimuli were presented in a pseudo-randomized order, with the constraint that no more than two stimuli of the same intensity occurred consecutively. To prevent tissue damage, the stimulation site was shifted by a few millimeters after each trial. After 40 trials, a short break was provided, during which participants were allowed to open their eyes and adjust their seating position.

### EEG recordings

Brain activity was recorded using 64 actiCAP slim/snap sensors placed according to the extended international 10-20 system (Easycap, Wörthsee, Germany) and BrainAmp MR plus amplifiers (Brain Products GmbH, Gilching, Germany). During recordings, sensors were referenced to FCz and grounded at Fpz. Signals were sampled with a sampling frequency of 1000 Hz and band-pass-filtered between 0.016 and 250 Hz. Impedances were kept below 20 kΩ.

### External replication data

To assess the replicability of results, we additionally included data from a large openly available database from Beijing, China (https://doi.org/10.18112/openneuro.ds005280.v1.0.0). From this database, we included the dataset associated with experiment no. 8 as it was both the largest available sample and most methodologically similar to our own Set_Munich in terms of stimulation site and EEG recording device. We refer to this dataset as Set_Beijing. The corresponding study protocol was approved by the ethics review committee of the Institute of Psychology, Chinese Academy of Sciences, and conducted in accordance with the Declaration of Helsinki. All participants provided written informed consent, including explicit consent for public data sharing.

The selected dataset included n = 223 healthy adults (93 male, 130 female; mean age = 20.8 ± 2.3 years). Recruitment procedures targeted healthy volunteers without known neurological or psychiatric disorders, neurosurgical history, or substance abuse. Participants were instructed to maintain a regular sleep–wake cycle and avoid alcohol and excessive caffeine consumption for at least 24 hours before testing. After preprocessing and exclusion of participants with insufficient artifact-free trials (see below), 111 participants remained for analysis. Similar to Set_Munich, this sample represents a specific, in this case, university-aged population and is therefore not representative of the general population in China or any other region.

Participants completed a single experimental session conducted in Mandarin Chinese. Prior to EEG setup, participants were familiarized with the nociceptive stimulation procedure. The following experimental paradigm consisted of 30 brief laser stimuli delivered to the dorsum of the left hand across 3 blocks of 10 trials. Stimuli were administered at two individually-calibrated intensity levels at 3.0 and 3.5 J or at 3.5 and 4.0 J with interstimulus intervals of 4–6 s. To be consistent with the trial selection performed in the analysis of Set_Munich, we only included trials with stimuli of the higher intensity for each individual. After each stimulus, participants verbally rated pain intensity on a numerical rating scale with 0 representing “no sensation”, 4 representing “onset of pain” (pinprick pain threshold), and 10 representing “worst pain imaginable”. Participants sat comfortably throughout the recording and wore protective goggles.

Nociceptive stimuli were delivered using a Nd:YAP laser system (1.34 µm wavelength, 4 ms pulse duration; spot diameter ∼7 mm), a device and stimulation protocol closely aligned with Set_Munich. Stimulation sites were slightly shifted after each trial to minimize local sensitization or tissue fatigue.

EEG data were acquired using a 64-channel Brain Products system (Brain Products GmbH, Germany) with Ag/AgCl electrodes positioned according to the international 10–20 system. Signals were recorded at a sampling frequency of 1000 Hz, referenced to the tip of the nose, and impedances were maintained below 10 kΩ.

### EEG preprocessing

BIDS-transformed [37] EEG data were automatically preprocessed using the automatic preprocessing part of the DISCOVER-EEG pipeline (v.2.0.0.; https://github.com/crisglav/discover-eeg) [38, 39] based on the MATLAB (R2020b, Mathworks, Natick, MA) toolbox EEGLAB (v2022.0) [40]. Please note that all analyses following automatic preprocessing were performed with additional code outside of DISCOVER-EEG (see below).

Automatic preprocessing included line noise removal, bad channel detection and interpolation, re-referencing to the average reference, independent component analysis (ICA), and the automatic detection of bad segments using the pipeline’s standard parameters. For Set_Beijing, the standard preprocessing procedure did not reliably detect and remove channels with large rhythmic signal discontinuities. Therefore, when preprocessing this dataset, we added an additional automated criterion for excluding artifactual channels: Channels were removed if their signal distribution contained more than one pronounced mode.

All 2.5-second peri-stimulus intervals (1.25 s before to 1.25 s after application of the laser stimulation) were concatenated prior to performing ICA and bad segment detection. Participants were included in subsequent analyses if at least five artifact-free trials remained at each of the included stimulus intensities. For the included participants of Set_Munich, the mean number of clean trials with stimuli of the two highest intensities ± SD (range) was 34.4 ± 6 (12-40) for session 1 and 35.2 ± 6 (12-40) for session 2. For the included participants of Set_Beijing, the mean number of clean trials with stimuli of the higher intensity ± SD (range) was 13.8 ± 2 (5-15).

### Extracting EEG responses

#### Individual-level responses

To quantify the neural response to painful stimuli, we quantified the canonical sequence of evoked potentials referred to as N1, N2, and P2 [41, 42] as well as induced neuronal oscillations at alpha (7 to <13 Hz), beta (14 to 30 Hz) and gamma (70 to 90 Hz) frequencies [8, 43–45]. Both single-trial evoked and oscillatory brain responses to noxious stimuli were quantified using established procedures [8, 46], using the MATLAB-based (R2021b, Mathworks, Natick, MA) toolbox FieldTrip (version: 20221022, [47]) along with custom-written code. To compute *evoked brain response amplitudes*, preprocessed data were bandpass-filtered between 1 and 30 Hz and baseline-corrected using the one-second pre-stimulus interval. Individual peak latencies of evoked responses were then determined based on averages across all trials. To this end, local minima/maxima of the averaged waveform were determined at predefined sensors (N1: C4, N2: Cz; P2: Cz) [8, 46, 48] and in predefined time-windows (N1: 120-200 ms; N2: 180-300 ms; P2: 250-500 ms) [8, 48]. Next, individual amplitudes were obtained by averaging across a 30-ms window [46, 48] centred at the previously defined peak latency. If no individual minimum or maximum could be detected, the amplitude at the midpoint of the corresponding pre-defined search window was used instead. To quantify the N1 response, data were additionally re-referenced to Fz [49] before computing the average waveform across all trials. To examine *oscillatory brain responses*, preprocessed data were filtered (1 Hz high-pass filter, 49 to 51 Hz band-stop filter). Single-trial time-frequency estimates were obtained using a Hanning-tapered Fast Fourier transformation and a sliding window approach, and subsequently averaged across trials. To obtain alpha and beta responses, a sliding window with a length of 500 ms and a step size of 20 ms was used. To obtain gamma responses, the window length was shortened to 250 ms, while the step size remained 20 ms. Finally, individual oscillatory responses were assessed by calculating the mean power within response-specific time-frequency-sensor windows (alpha: 7-12 Hz, 500-900 ms, averaged across Cz, CPz, C2, C4, CP2, CP4 [8, 44, 48]; beta: 14-30 Hz, 300-600 ms, averaged across Cz, CPz, C2, C4, CP2, CP4 [8, 44, 48]; gamma: 70-90 Hz, 150-350 ms, Cz [8, 46]). The procedure for computing individual-level responses was applied independently to the internal session one, internal session two, and the external dataset.

#### Trial-level responses

Trial-level evoked responses were quantified using the same processing, sensors, and peak detection procedures as described for the individual-level responses. Single-trial amplitudes were computed by averaging the signal for each trial within a 30-ms window centred at each participant’s previously defined individual peak latency. Trial-level oscillatory responses were also quantified using the same processing and response-specific time-frequency-sensor windows as described for the individual-level responses. Single-trial responses were computed by averaging the power within the respective time-frequency-sensor windows for each trial separately. The procedure for computing trial-level responses was applied independently to the internal session one, internal session two, and the external dataset.

### Statistical analyses

#### Models of inter-individual variability

To investigate the associations between EEG responses and inter-individual variability of pain perception, we performed Bayesian multi-model linear regression. We implemented the analysis in the R programming environment [50] using the “BayesFactor” package [51]. Our implementation’s results are consistent with those generated by the corresponding functionality of the JASP software (version: 0.19.3, [52]). In our models, individual-level brain responses (N1, N2, P2, alpha, beta, gamma) were the predictors, and individual-level pain ratings were the outcome variable. To obtain individual-level pain ratings, trial-level pain ratings were first averaged per intensity and then per subject to ensure equal weighting of each intensity in the analysis. To make predictor coefficients more comparable, we performed Z-standardization of all predictor and outcome variables across all individuals. Since Bayesian multi-model linear regression assumes linear relationships between the variables, scatter plots showing the relations between the different predictor variables and pain rating were visually inspected [20]. All relations appeared approximately linear by eye. Thus, no variable transformation was performed, and no variables needed to be excluded from further analyses. As recommended for Bayesian multi-model linear regression [20], analyses used the default settings also implemented in JASP [52]. Thus, as the prior on parameters, we used a Jeffreys-Zellner-Siow (JZS) r-scale prior with a width of 0.354. As the prior on the models, we employed a binomial model prior with alpha = beta = 1. Statistical inferences were based on Bayes factors (BF; see below). For model selection, the ratio of prior and posterior model odds (BF_M_) as well as the evidence of a given model relative to the null model (BF_10_) were assessed. In order to evaluate the relative importance of individual predictors, coefficient estimates as well as posterior inclusion probabilities (BF_inclusion_), i.e., the factor by which the odds of including a predictor have changed after observing the data, averaged across all models that include or exclude that predictor, were obtained using Bayesian model averaging [20].

#### Models of intra-individual variability

To investigate the associations between EEG responses and intra-individual variability of pain perception at the moment-to-moment timescale, we performed Bayesian linear mixed-effects modelling using custom-written R code [50] in RStudio (version: 2024.12.0.467; [53]), based on the “BayesFactor” package [51]. This implementation closely followed the model structure and priors used for the inter-individual analysis and implemented in the JASP software [52], but extended it to accommodate random effects. Specifically, we introduced a random (i.e., subject-specific) intercept term to account for the repeated measures design. The outcome variable was the single-trial pain rating, and predictors were single-trial brain responses (N1, N2, P2, alpha, beta, gamma). To eliminate stimulus intensity-related variability, single-trial pain ratings and brain responses were mean-centred per subject and intensity. To make predictor coefficients more comparable, we performed Z-standardization of all predictor and outcome variables across all individuals and trials. Bayesian inference was applied in the same way as in the analysis of inter-individual variability, including the specification of priors, model selection procedures, the use of model-averaged posterior estimates, and inclusion probabilities to evaluate predictor relevance.

To assess the associations between EEG responses and intra-individual variability of pain perception at the day-to-day timescale, we proceeded analogously. However, instead of single-trial data, we analyzed the individual-level data concatenated across the two internal recording sessions. This resulted in two datapoints per individual of which we subtracted the mean per individual to remove any inter-individual variability.

#### Inference criteria

All inferences were based on Bayes factors following [54]. Each model was evaluated using three metrics: First, BF_M_ quantifies the change from prior to posterior model odds after observing the data, relative to all other models considered. Evidence for increased (decreased) model odds was inferred when BF_M_ > 3 (or < 1/3). Second, BF_10_ quantifies the adequacy of a model in explaining the data relative to the null model. Evidence for (or against) the model relative to the null was inferred when BF_10_ > 3 (or < 1/3). In all analyses, the null model included a fixed (i.e., not subject-specific) intercept term. In analyses of inter-individual variability, the null model additionally included age and sex covariates. In analyses of intra-individual variability, the null model did not comprise the random intercept, since all inter-individual variability had been removed through within-individual centring of the data prior to model fitting. Third, R^2^ represents the explained variance of a model and is therefore a measure of the accuracy of model predictions. Importantly, since Bayes Factors impose a penalty for model complexity, the highest BF_10_ is not necessarily the one with the largest R^2^. Finally, the evidence for including individual predictors was assessed using BF_inclusion_, which reflects the change from prior to posterior inclusion odds averaged across all models. Evidence for (or against) the inclusion of a predictor was inferred when BF_inclusion_ > 3 (or < 1/3).

#### Robustness

We evaluated the robustness of our findings by testing both their repeatability and replicability. To assess repeatability, defined as obtaining consistent results when repeating the experiment in the same cohort, we tested whether the model-averaged mean predictor coefficients estimated from the first internal recording session remained informative in the second internal recording session, and vice versa. Specifically, we predicted pain ratings in the second session using the second session’s brain responses together with the model-averaged coefficients from the first session. We then quantified the association between predicted and true pain ratings using a Bayesian correlation analysis for inter-individual models, and a Bayesian linear mixed effects model with a random (i.e., subject-specific) intercept term for intra-individual models. To assess replicability, defined as obtaining consistent results when performing a similar experiment in a different cohort, we proceeded analogously, except that the datasets used for coefficient estimation and testing came from distinct participant cohorts.

#### Statistical power

Since our analyses are based on Bayesian multi-model regression and Bayesian linear mixed-effects models (LMEs), which are not currently supported by Bayes factor design analysis (BFDA)[55] tools, we performed an approximate power analysis using a Bayesian correlation model as a proxy. Using the R package “BFDA” [56] with default priors and 10,000 simulations, we estimated statistical power for our sample size of n = 161 participants. Assuming a medium-sized correlation (R = 0.3) and BF_10_ thresholds of 1/3 and 3, the analysis yielded a statistical power of 94% with a false positive rate of 0.2%. Under more conservative thresholds (BF_10_ = 1/10 and 10), power was 86% with 0% false positives. These simulations serve as an approximate indication that the study was well-powered to detect medium-sized effects. While they do not fully reflect the complexity of the models, i.e. their hierarchical structure, they offer a reasonable estimate in the absence of validated Bayesian power tools for regression and mixed-effects models.

## Data availability

Raw and preprocessed data in standardized EEG-BIDS format [37] and the code used for the current manuscript will be made available upon acceptance of the manuscript or request by the reviewing editor.

## Acknowledgments

The study was supported by the Deutsche Forschungsgemeinschaft (PL 321/14-1, PL321/16-1, SFB1158) and the Technical University of Munich (TUM Innovation Network Neurotech). We thank Li Hu for sharing data.

## Author contributions

Laura Tiemann: Conceptualization, Methodology, Software, Investigation, Data Curation, Writing – Original Draft, Writing – Review & Editing

Felix S. Bott: Conceptualization, Methodology, Software, Formal analysis, Validation, Data Curation, Writing – Original Draft, Visualization, Writing-Review & Editing

Elisabeth S. May: Conceptualization, Writing – Review & Editing

Moritz M. Nickel: Conceptualization, Writing – Review & Editing

Vanessa D. Hohn: Conceptualization

Cristina Gil Ávila: Conceptualization

Nicolò Bruna: Writing – Review & Editing

Paul Theo Zebhauser: Conceptualization, Writing – Review & Editing

Markus Ploner: Conceptualization, Resources, Writing – Original Draft, Writing – Review & Editing, Visualization, Supervision, Project Administration, Funding acquisition

## Supplementary Materials

**Figure S1:**
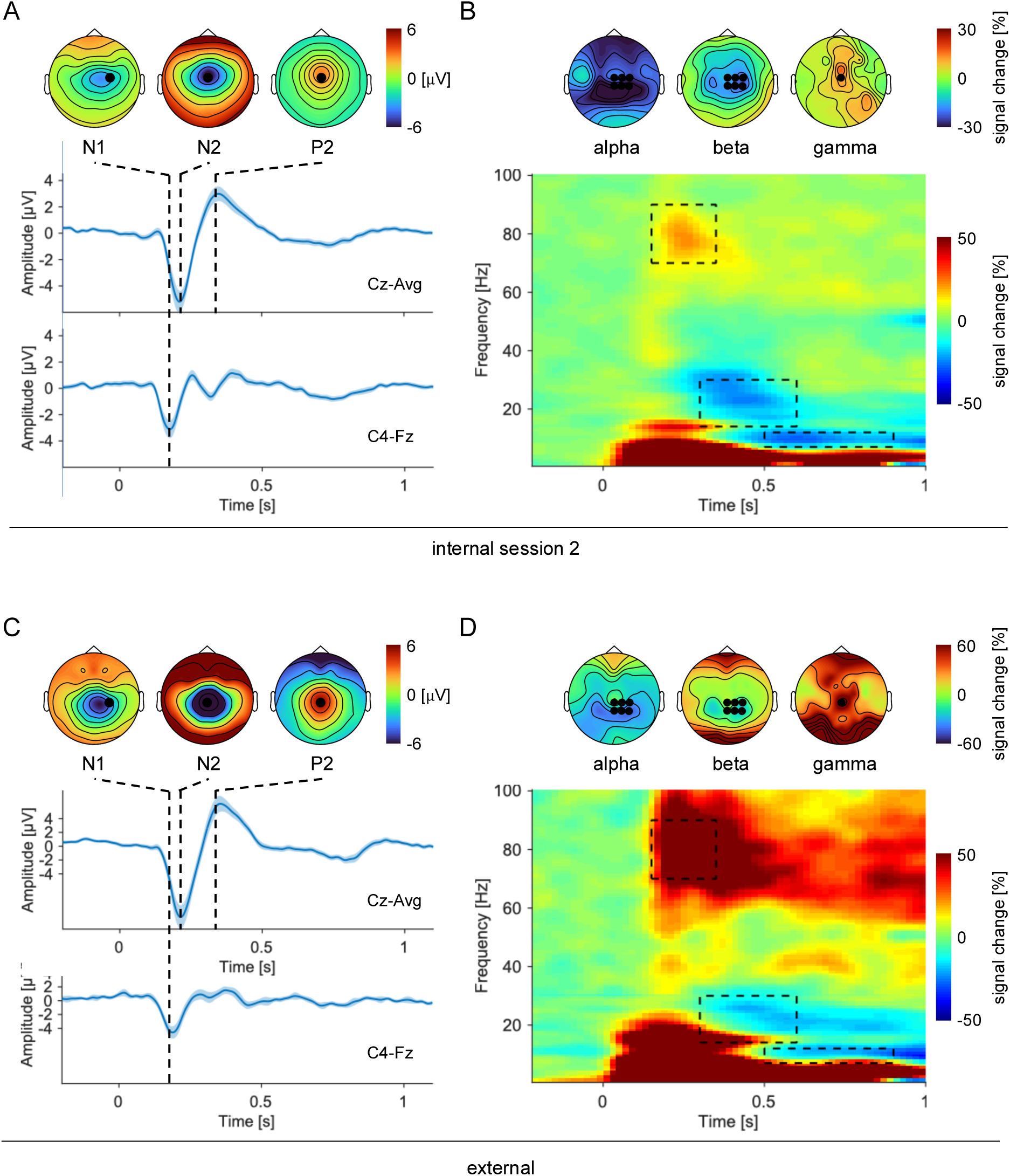
Brain responses to noxious stimuli in the internal session 2 (A + B) and external data (C + D). A) + C) Grand average of event-related potentials for the Cz-Avg montage (to visualize the N2 and P2 responses) and the C4-Fz montage (to visualize the N1 response). The shaded blue bands represent +/- 2 SEM. Scalp topographies show the spatial distributions of voltage for the respective montages. \ Positions of measurement electrodes are marked as black dots. B) + D) Grand average time-frequency representation (TFR) showing changes in oscillatory power relative to a pre-stimulus baseline [-1 to 0 s]. For visualization, TFRs at Cz are presented. Scalp topographies illustrate the spatial distributions of relative power changes, averaged within the indicated time-frequency windows. Measurement electrode included later for the quantification of oscillatory power in the different frequency bands are marked as black dots.

**Figure S2:**
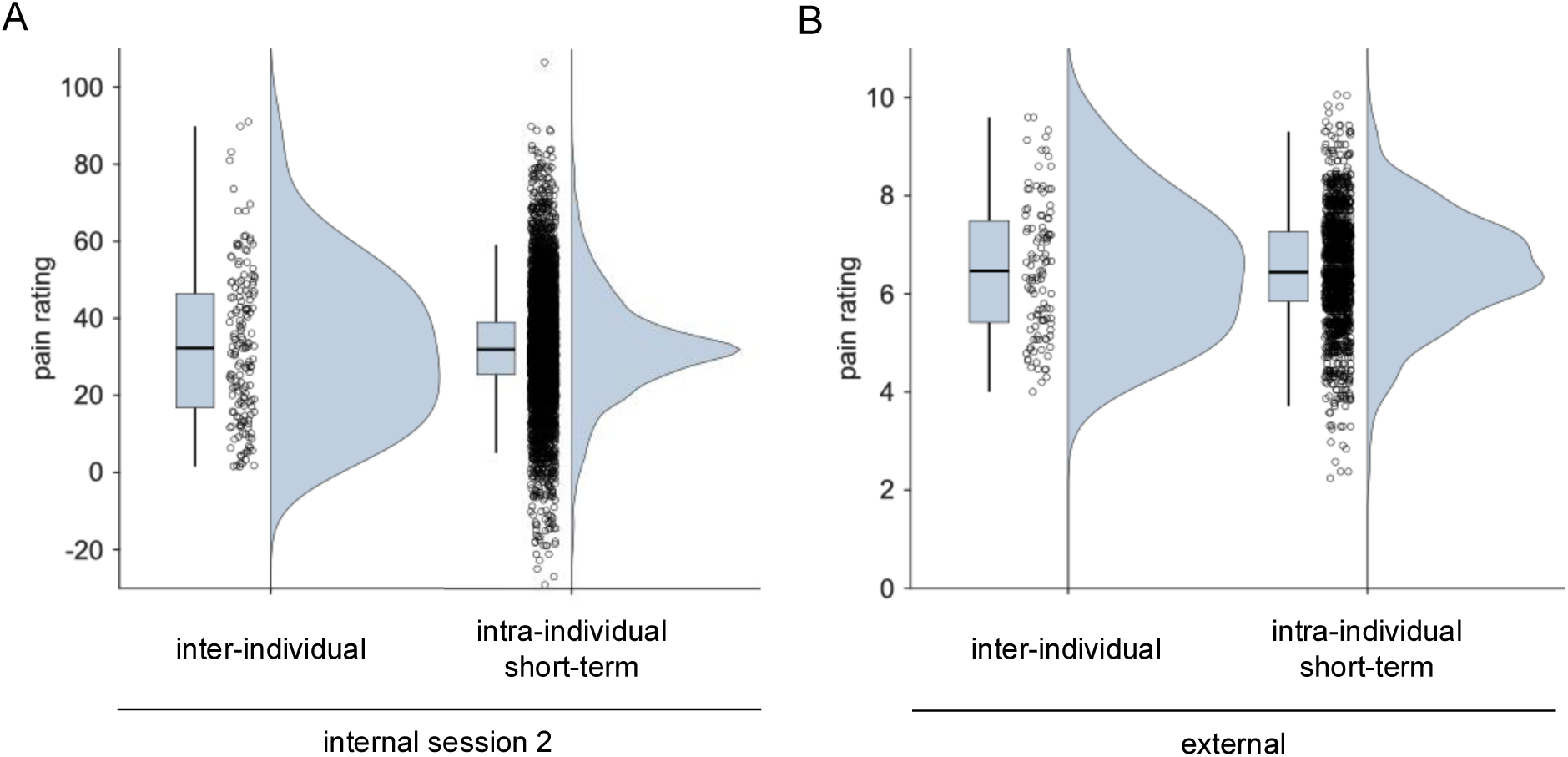
Pain ratings in the internal session 2 (A) and external data (B). A) + B) Left: Individual pain ratings reflecting inter-individual pain variability were obtained by averaging single-trail pain ratings for each individual, first within each stimulus intensity and then across stimulus intensities, yielding one rating value per individual. Right: Trial-level pain ratings (left) reflecting intra-individual short-term pain variability were obtained by computing the means of single-trial pain ratings for each individual and stimulus intensity, and then subtracting these means from the corresponding single-trail pain ratings, yielding one centred rating value per trial. For visualization only, trial-wise and session-wise pain ratings were re-centred to the mean of the inter-individual distribution.

**Figure S3:**
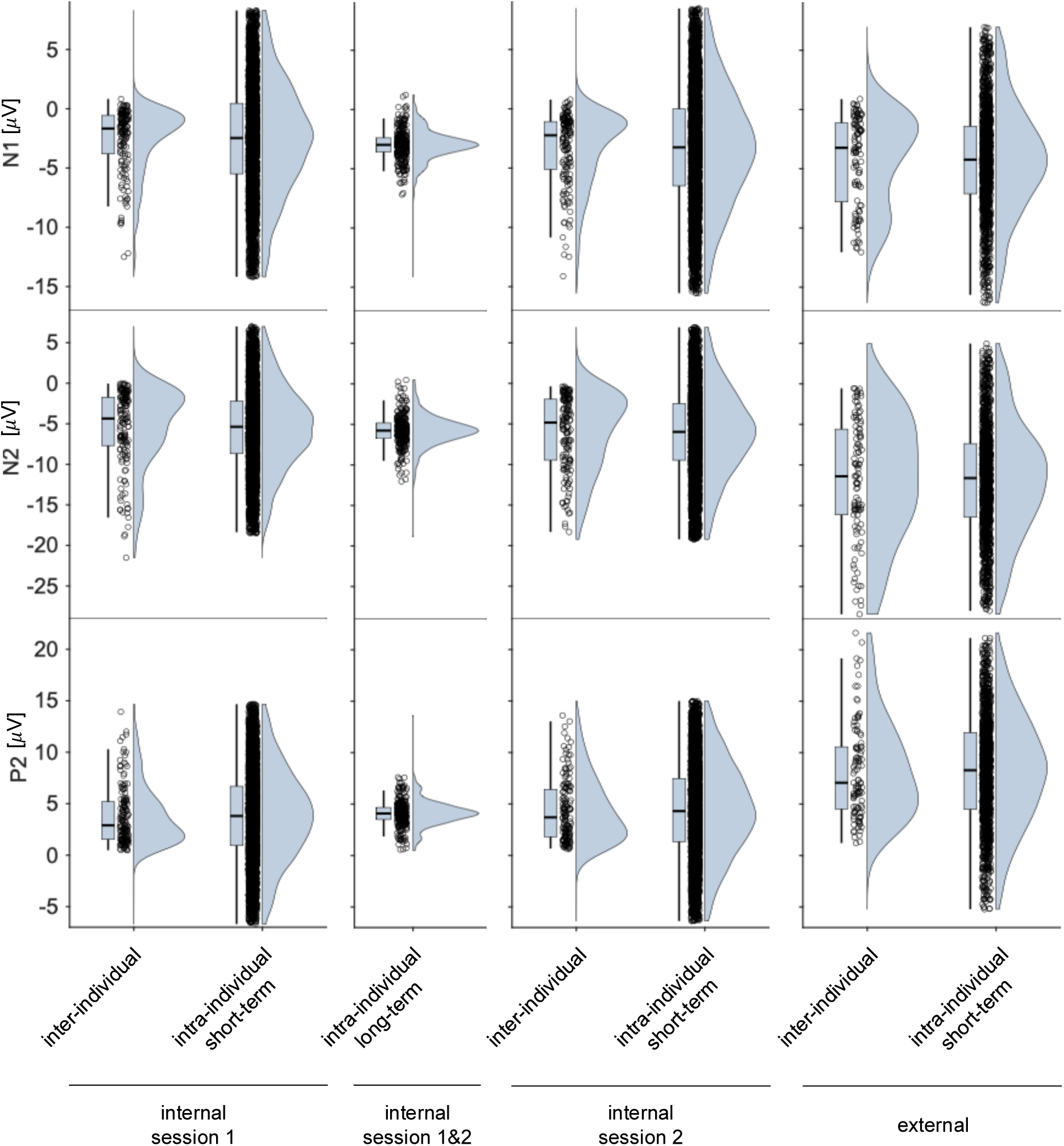
Evoked potential brain responses. To enhance interpretability, the lowest and highest 2.5 percent of data points are excluded from each raincloud plot. Otherwise, the visualization of brain response distributions is analogous to the visualization of pain ratings in Figures 3 and S2.

**Figure S4:**
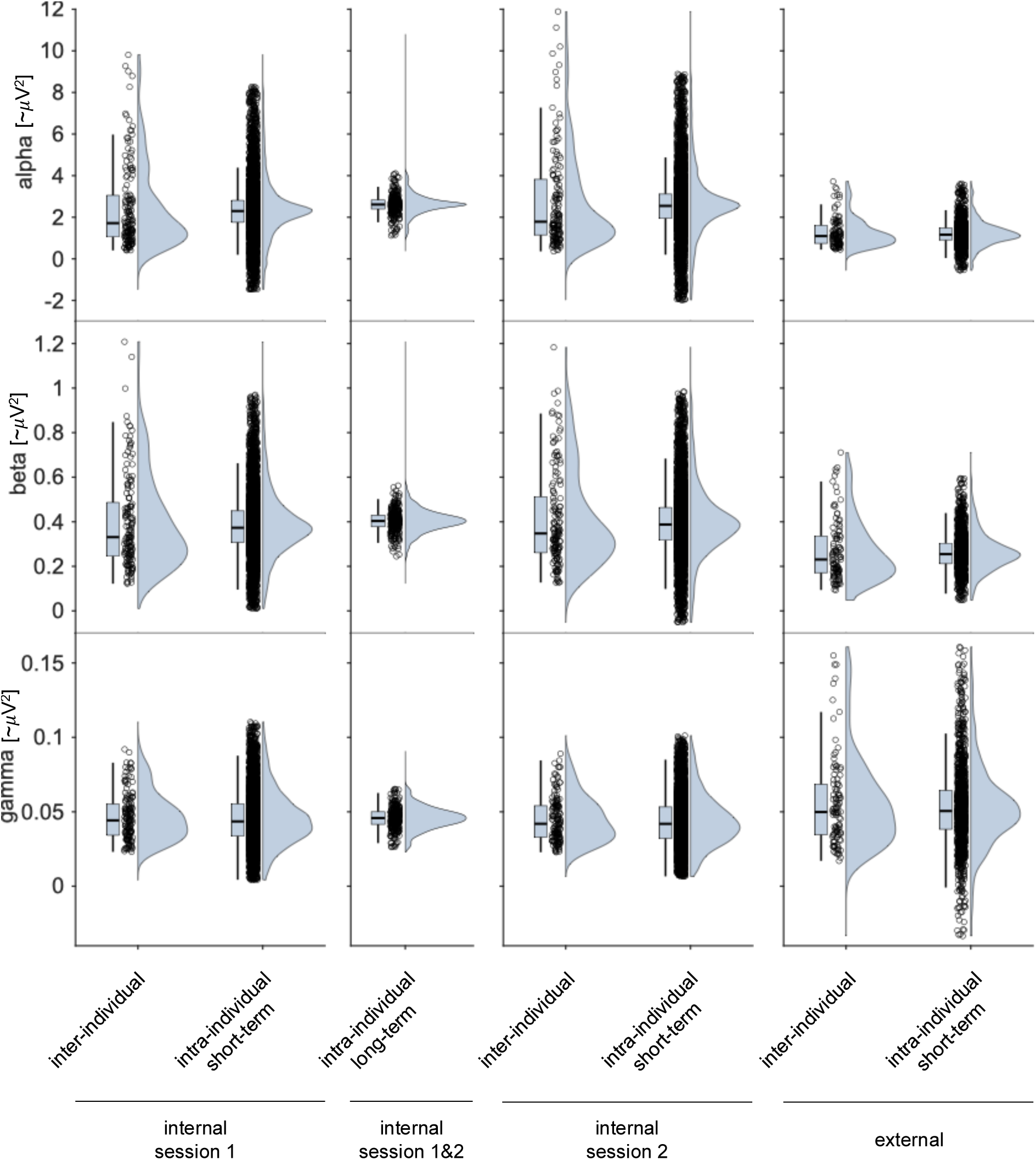
Induced oscillatory brain responses. To enhance interpretability, the lowest and highest 2.5 percent of data points are excluded from each raincloud plot. Otherwise, the visualization of brain response distributions is analogous to the visualization of pain ratings in Figures 3 and S2.

**Figure S5:**
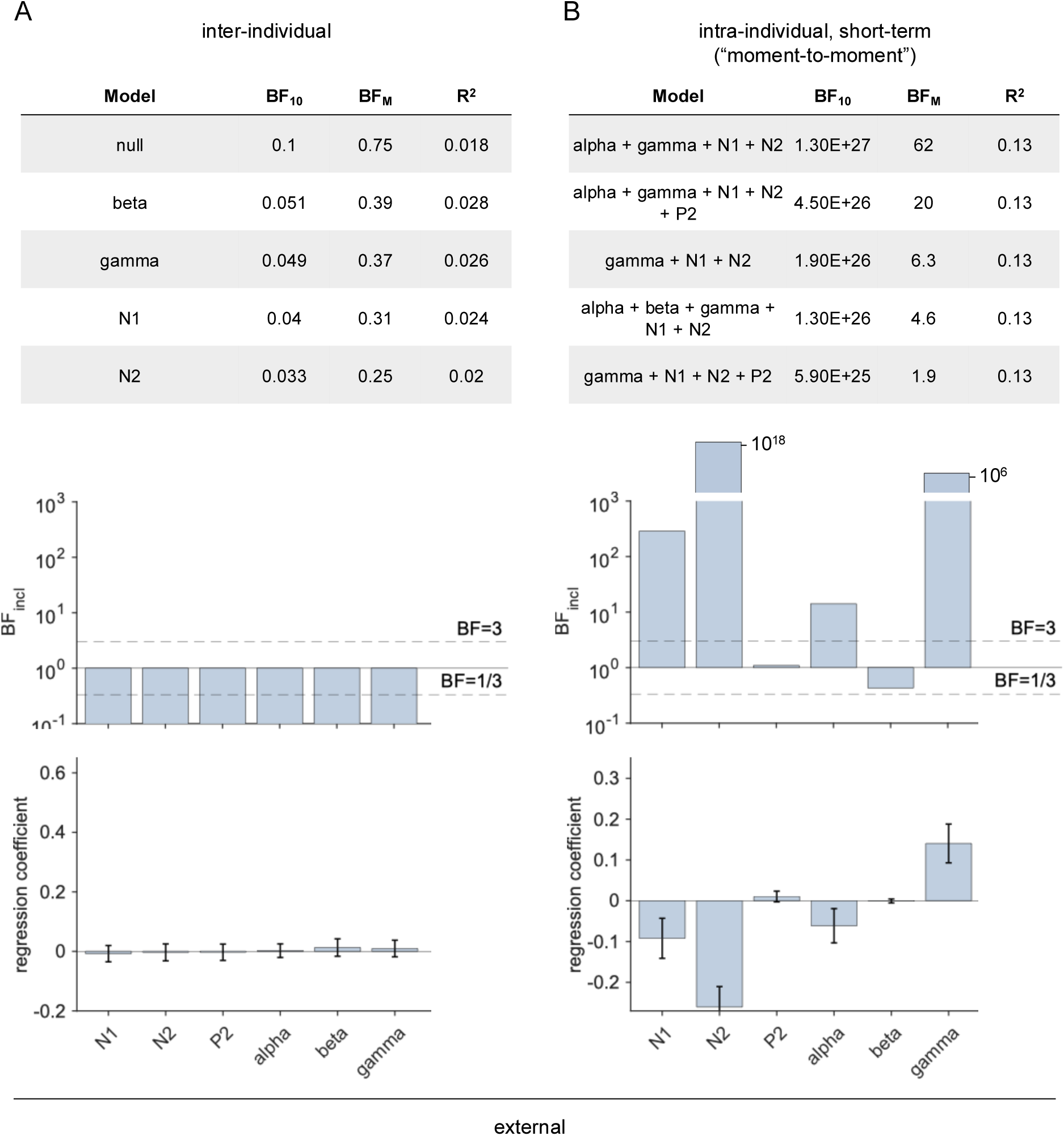
Bayesian multi-model analysis of inter-individual (A) intra-individual (B) variability in the external data. A) + B) Top: The five models with the highest evidence relative to the null model (BF_10_) are shown, along with their ratios of prior and posterior model odds (BF_M_) and their explained variance (R^2^). Middle: Inclusion Bayes Factors (BF_incl_), representing the evidence for including each predictor averaged across all models considered. Bottom: Model-averaged posterior mean estimates for each predictor coefficient. Error bars indicate +/-2 standard deviations of the model-averaged posterior coefficient distribution.

## References

1. Mun CJ, Suk HW, Davis MC, Karoly P, Finan P, Tennen H, et al. Investigating intraindividual pain variability: methods, applications, issues, and directions. Pain. 2019;160(11):2415–29. Epub 2019/05/31. doi: 10.1097/j.pain.0000000000001626. PubMed PMID: 31145212.

2. Nielsen CS, Staud R, Price DD. Individual differences in pain sensitivity: measurement, causation, and consequences. The journal of pain : official journal of the American Pain Society. 2009;10(3):231–7. PubMed PMID: 19185545.

3. Ossipov MH, Dussor GO, Porreca F. Central modulation of pain. The Journal of clinical investigation. 2010;120(11):3779–87. Epub 20101101. doi: 10.1172/JCI43766. PubMed PMID: 21041960; PubMed Central PMCID: PMCPMC2964993.

4. Madden VJ, Kamerman PR, Catley MJ, Bellan V, Russek LN, Camfferman D, et al. Variability in experimental pain studies: nuisance or opportunity? Br J Anaesth. 2021;126(2):e61–e4. Epub 20201217. doi: 10.1016/j.bja.2020.11.005. PubMed PMID: 33341221; PubMed Central PMCID: PMCPMC8014940.

5. Gim S, Lee DH, Lee S, Woo CW. Interindividual differences in pain can be explained by fMRI, sociodemographic, and psychological factors. Nature communications. 2024;15(1):7883. Epub 20240910. doi: 10.1038/s41467-024-51910-9. PubMed PMID: 39256362; PubMed Central PMCID: PMCPMC11387422.

6. Hoeppli ME, Nahman-Averbuch H, Hinkle WA, Leon E, Peugh J, Lopez-Sola M, et al. Dissociation between individual differences in self-reported pain intensity and underlying fMRI brain activation. Nature communications. 2022;13(1):3569. Epub 20220622. doi: 10.1038/s41467-022-31039-3. PubMed PMID: 35732637.

7. Hoeppli ME, Nahman-Averbuch H, Hinkle WA, Leon E, Peugh J, Lopez-Sola M, et al. Reply to: Interindividual differences in pain can be explained by fMRI, sociodemographic, and psychological factors. Nature communications. 2024;15(1):7884. Epub 20240910. doi: 10.1038/s41467-024-51911-8. PubMed PMID: 39256345; PubMed Central PMCID: PMCPMC11387405.

8. Hu L, Iannetti GD. Neural indicators of perceptual variability of pain across species. Proceedings of the National Academy of Sciences of the United States of America. 2019. doi: 10.1073/pnas.1812499116. PubMed PMID: 30642968.

9. Schulz E, Tiemann L, Schuster T, Gross J, Ploner M. Neurophysiological coding of traits and states in the perception of pain. Cereb Cortex. 2011;21:2408–14.

10. Schulz E, Zherdin A, Tiemann L, Plant C, Ploner M. Decoding an Individual’s Sensitivity to Pain from the Multivariate Analysis of EEG Data. Cereb Cortex. 2012;22:1118–23. doi: 10.1093/cercor/bhr186. PubMed PMID: 21765182.

11. Kotikalapudi R, Kincses B, Zunhammer M, Schlitt F, Asan L, Schmidt-Wilcke T, et al. Brain morphology predicts individual sensitivity to pain: a multicenter machine learning approach. Pain. 2023;164(11):2516–27. Epub 2023/06/15. doi: 10.1097/j.pain.0000000000002958. PubMed PMID: 37318027; PubMed Central PMCID: PMCPMC10578427 interests that may be relevant to content are disclosed at the end of this article.

12. Spisak T, Kincses B, Schlitt F, Zunhammer M, Schmidt-Wilcke T, Kincses ZT, et al. Pain-free resting-state functional brain connectivity predicts individual pain sensitivity. Nature communications. 2020;11(1):187. Epub 2020/01/12. doi: 10.1038/s41467-019-13785-z. PubMed PMID: 31924769; PubMed Central PMCID: PMCPMC6954277.

13. Coghill RC, Sang CN, Maisog JM, Iadarola MJ. Pain intensity processing within the human brain: a bilateral, distributed mechanism. J Neurophysiol. 1999;82(4):1934–43.

14. Huang G, Xiao P, Hung YS, Iannetti GD, Zhang ZG, Hu L. A novel approach to predict subjective pain perception from single-trial laser-evoked potentials. NeuroImage. 2013;81:283–93. Epub 2013/05/21. doi: 10.1016/j.neuroimage.2013.05.017. PubMed PMID: 23684861.

15. Li L, Huang G, Lin Q, Liu J, Zhang S, Zhang Z. Magnitude and Temporal Variability of Inter-stimulus EEG Modulate the Linear Relationship Between Laser-Evoked Potentials and Fast-Pain Perception. Front Neurosci. 2018;12:340. Epub 20180531. doi: 10.3389/fnins.2018.00340. PubMed PMID: 29904336; PubMed Central PMCID: PMCPMC5991169.

16. Mayhew SD, Hylands-White N, Porcaro C, Derbyshire SW, Bagshaw AP. Intrinsic variability in the human response to pain is assembled from multiple, dynamic brain processes. NeuroImage. 2013;75:68–78. Epub 2013/03/15. doi: 10.1016/j.neuroimage.2013.02.028. PubMed PMID: 23485593.

17. Garcia-Larrea L, Peyron R, Laurent B, Mauguiere F. Association and dissociation between laser-evoked potentials and pain perception. Neuroreport. 1997;8(17):3785–9. PubMed PMID: 9427371.

18. Iannetti GD, Zambreanu L, Cruccu G, Tracey I. Operculoinsular cortex encodes pain intensity at the earliest stages of cortical processing as indicated by amplitude of laser-evoked potentials in humans. Neuroscience. 2005;131(1):199–208. doi: 10.1016/j.neuroscience.2004.10.035. PubMed PMID: 15680703.

19. Gross J, Schnitzler A, Timmermann L, Ploner M. Gamma oscillations in human primary somatosensory cortex reflect pain perception. PLoS Biol. 2007;5(5):e133. Epub 2007/04/26. doi: 10.1371/journal.pbio.0050133. PubMed PMID: 17456008; PubMed Central PMCID: PMC1854914.

20. Bergh DVD, Clyde MA, Gupta A, de Jong T, Gronau QF, Marsman M, et al. A tutorial on Bayesian multi-model linear regression with BAS and JASP. Behavior research methods. 2021;53(6):2351–71. Epub 20210409. doi: 10.3758/s13428-021-01552-2. PubMed PMID: 33835394; PubMed Central PMCID: PMCPMC8613115.

21. Nickel MM, Tiemann L, Hohn VD, May ES, Gil Avila C, Eippert F, et al. Temporal-spectral signaling of sensory information and expectations in the cerebral processing of pain. Proceedings of the National Academy of Sciences of the United States of America. 2022;119(1). doi: 10.1073/pnas.2116616119. PubMed PMID: 34983852; PubMed Central PMCID: PMCPMC8740684.

22. Bott FS, Nickel MM, Hohn VD, May ES, Gil Avila C, Tiemann L, et al. Local brain oscillations and interregional connectivity differentially serve sensory and expectation effects on pain. Sci Adv. 2023;9(16):eadd7572. Epub 2023/04/19. doi: 10.1126/sciadv.add7572. PubMed PMID: 37075123; PubMed Central PMCID: PMCPMC10115421.

23. Kohoutova L, Atlas LY, Buchel C, Buhle JT, Geuter S, Jepma M, et al. Individual variability in brain representations of pain. Nat Neurosci. 2022;25(6):749–59. Epub 2022/06/01. doi: 10.1038/s41593-022-01081-x. PubMed PMID: 35637368; PubMed Central PMCID: PMCPMC9435464.

24. Zhang LB, Geng XY, Hu L. Neural variability reliably encodes interindividual differences in the perception of pain intensity. PLoS Biol. 2025;23(10):e3003470. Epub 2025/10/27. doi: 10.1371/journal.pbio.3003470. PubMed PMID: 41144553; PubMed Central PMCID: PMCPMC12574952.

25. Barroso J, Branco P, Apkarian AV. Brain mechanisms of chronic pain: critical role of translational approach. Transl Res. 2021;238:76–89. Epub 2021/06/29. doi: 10.1016/j.trsl.2021.06.004. PubMed PMID: 34182187; PubMed Central PMCID: PMCPMC8572168.

26. Reddan MC. Recommendations for the Development of Socioeconomically-Situated and Clinically-Relevant Neuroimaging Models of Pain. Frontiers in neurology. 2021;12:700833. Epub 2021/09/25. doi: 10.3389/fneur.2021.700833. PubMed PMID: 34557144; PubMed Central PMCID: PMCPMC8453079.

27. Woo CW, Chang LJ, Lindquist MA, Wager TD. Building better biomarkers: brain models in translational neuroimaging. Nat Neurosci. 2017;20(3):365–77. doi: 10.1038/nn.4478. PubMed PMID: 28230847.

28. May ES, Tiemann L, Gil Avila C, Bott FS, Hohn VD, Gross J, et al. Assessing the predictive value of peak alpha frequency for the sensitivity to pain. Pain. 2025. Epub 20250311. doi: 10.1097/j.pain.0000000000003571. PubMed PMID: 40085759.

29. Ozcan A, Tulum Z, Pinar L, Baskurt F. Comparison of pressure pain threshold, grip strength,dexterity and touch pressure of dominant and non-dominant hands within and between right-and left-handed subjects. J Korean Med Sci. 2004;19(6):874–8. doi: 10.3346/jkms.2004.19.6.874. PubMed PMID: 15608401; PubMed Central PMCID: PMCPMC2816288.

30. Pud D, Golan Y, Pesta R. Hand dominancy--a feature affecting sensitivity to pain. Neurosci Lett. 2009;467(3):237–40. Epub 20091021. doi: 10.1016/j.neulet.2009.10.048. PubMed PMID: 19853018.

31. Epperson CN, Haga K, Mason GF, Sellers E, Gueorguieva R, Zhang W, et al. Cortical gamma-aminobutyric acid levels across the menstrual cycle in healthy women and those with premenstrual dysphoric disorder: a proton magnetic resonance spectroscopy study. Arch Gen Psychiatry. 2002;59(9):851–8. doi: 10.1001/archpsyc.59.9.851. PubMed PMID: 12215085.

32. Iacovides S, Avidon I, Baker FC. Does pain vary across the menstrual cycle? A review. European journal of pain (London, England). 2015;19(10):1389–405. Epub 20150421. doi: 10.1002/ejp.714. PubMed PMID: 25899177.

33. Sumner RL, McMillan RL, Shaw AD, Singh KD, Sundram F, Muthukumaraswamy SD. Peak visual gamma frequency is modified across the healthy menstrual cycle. Hum Brain Mapp. 2018;39(8):3187–202. Epub 2018/04/18. doi: 10.1002/hbm.24069. PubMed PMID: 29665216; PubMed Central PMCID: PMCPMC6055613.

34. Tan HM, Gross J, Uhlhaas PJ. MEG sensor and source measures of visually induced gamma-band oscillations are highly reliable. NeuroImage. 2016;137:34–44. Epub 20160503. doi: 10.1016/j.neuroimage.2016.05.006. PubMed PMID: 27153980; PubMed Central PMCID: PMCPMC5405052.

35. Hu L, Zhang ZG, Mouraux A, Iannetti GD. Multiple linear regression to estimate time-frequency electrophysiological responses in single trials. NeuroImage. 2015;111:442–53. Epub 2015/02/11. doi: 10.1016/j.neuroimage.2015.01.062. PubMed PMID: 25665966; PubMed Central PMCID: PMCPMC4401443.

36. Adamczyk WM, Szikszay TM, Nahman-Averbuch H, Skalski J, Nastaj J, Gouverneur P, et al. To Calibrate or not to Calibrate? A Methodological Dilemma in Experimental Pain Research. The journal of pain : official journal of the American Pain Society. 2022;23(11):1823–32. Epub 20220731. doi: 10.1016/j.jpain.2022.07.007. PubMed PMID: 35918020.

37. Pernet CR, Appelhoff S, Gorgolewski KJ, Flandin G, Phillips C, Delorme A, et al. EEG-BIDS, an extension to the brain imaging data structure for electroencephalography. Sci Data. 2019;6(1):103. Epub 2019/06/27. doi: 10.1038/s41597-019-0104-8. PubMed PMID: 31239435; PubMed Central PMCID: PMCPMC6592877.

38. Gil Ávila C, Bott FS, Tiemann L, Hohn VD, May ES, Nickel MM, et al. DISCOVER-EEG: an open, fully automated EEG pipeline for biomarker discovery in clinical neuroscience. Sci Data. 2023;10(1):613. Epub 20230911. doi: 10.1038/s41597-023-02525-0. PubMed PMID: 37696851; PubMed Central PMCID: PMCPMC10495446.

39. Pernet CR, Martinez-Cancino R, Truong D, Makeig S, Delorme A. From BIDS-Formatted EEG Data to Sensor-Space Group Results: A Fully Reproducible Workflow With EEGLAB and LIMO EEG. Front Neurosci. 2020;14:610388. doi: 10.3389/fnins.2020.610388. PubMed PMID: 33519362; PubMed Central PMCID: PMCPMC7845738.

40. Delorme A, Makeig S. EEGLAB: an open source toolbox for analysis of single-trial EEG dynamics including independent component analysis. Journal of neuroscience methods. 2004;134(1):9–21. doi: 10.1016/j.jneumeth.2003.10.009. PubMed PMID: 15102499.

41. Garcia-Larrea L, Frot M, Valeriani M. Brain generators of laser-evoked potentials: from dipoles to functional significance. Neurophysiol Clin. 2003;33(6):279–92. PubMed PMID: 14678842.

42. Lorenz J, Garcia-Larrea L. Contribution of attentional and cognitive factors to laser evoked brain potentials. Neurophysiol Clin. 2003;33(6):293–301. PubMed PMID: 14678843.

43. Kim JA, Davis KD. Neural Oscillations: Understanding a Neural Code of Pain. Neuroscientist. 2020:1073858420958629. Epub 2020/09/29. doi: 10.1177/1073858420958629. PubMed PMID: 32981457.

44. Pernet CR, Garrido MI, Gramfort A, Maurits N, Michel CM, Pang E, et al. Issues and recommendations from the OHBM COBIDAS MEEG committee for reproducible EEG and MEG research. Nat Neurosci. 2020;23(12):1473–83. Epub 2020/09/23. doi: 10.1038/s41593-020-00709-0. PubMed PMID: 32958924.

45. Ploner M, Sorg C, Gross J. Brain Rhythms of Pain. Trends in cognitive sciences. 2017;21(2):100–10. doi: 10.1016/j.tics.2016.12.001. PubMed PMID: 28025007.

46. Tiemann L, Hohn VD, Ta Dinh S, May ES, Nickel MM, Gross J, et al. Distinct patterns of brain activity mediate perceptual and motor and autonomic responses to noxious stimuli. Nature communications. 2018;9(1):4487. Epub 2018/10/28. doi: 10.1038/s41467-018-06875-x. PubMed PMID: 30367033.

47. Oostenveld R, Fries P, Maris E, Schoffelen JM. FieldTrip: Open source software for advanced analysis of MEG, EEG, and invasive electrophysiological data. Computational intelligence and neuroscience. 2011;2011:156869. doi: 10.1155/2011/156869. PubMed PMID: 21253357; PubMed Central PMCID: PMC3021840.

48. Hohn VD, Bott FS, May ES, Tiemann L, Fritzen C, Nickel MM, et al. How do alpha oscillations shape the perception of pain? – An EEG-based neurofeedback study. OSF. 2022. doi: 10.17605/osf.io/qbkj2.

49. Hu L, Mouraux A, Hu Y, Iannetti GD. A novel approach for enhancing the signal-to-noise ratio and detecting automatically event-related potentials (ERPs) in single trials. NeuroImage. 2010;50(1):99–111. doi: 10.1016/j.neuroimage.2009.12.010. PubMed PMID: 20004255.

50. R Core Team. R: A Language and Environement for Statistical Computing. Vienna, Austria: R Foundation for Statistical Computing; 2021.

51. Morey RD, Rouder JN. BayesFactor: Computation of Bayes Factors for Common Designs. R package version 0.9.12–4.2. 2018.

52. JASP. JASP Team, 2025. (Version 0.19.3)[Computer software]. 2025.

53. Posit Team. RStudio: Integrated Development Environment for R. Boston, MA: Posit Software, PBC; 2025.

54. Jeffreys H. Theory of probability, 3rd edn oxford: Oxford university press. 1961.

55. Schönbrodt FD, Wagenmakers EJ. Bayes factor design analysis: Planning for compelling evidence. Psychon Bull Rev. 2018;25(1):128–42. Epub 2017/03/03. doi: 10.3758/s13423-017-1230-y. PubMed PMID: 28251595.

56. Schönbrodt FD, Stefan AM. BFDA: An R package for Bayes factor design analysis (version 0.5.0). 2019.

